# DDK regulates replication initiation by controlling the multiplicity of Cdc45-GINS binding to Mcm2-7

**DOI:** 10.1101/2020.12.09.417428

**Authors:** Lorraine De Jesús-Kim, Larry J. Friedman, Christian Ramsoomair, Jeff Gelles, Stephen P. Bell

**Author notes:** Co-corresponding authors: Stephen P. Bell, Jeff Gelles.

## Abstract

The committed step of eukaryotic DNA replication occurs when the replicative Mcm2-7 helicase pairs that license each replication origin are activated. Helicase activation requires the recruitment of Cdc45 and GINS to Mcm2-7, forming Cdc45-Mcm2-7-GINS complexes (CMGs). Using single-molecule biochemical assays to monitor CMG formation, we found that Cdc45 and GINS are recruited to loaded Mcm2-7 in two stages. Initially, Cdc45 and GINS are individually recruited to unstructured Mcm2-7 N-terminal tails in a Dbf4-dependent kinase (DDK)-dependent manner, forming Cdc45-tail-GINS intermediates (CtGs). The multiple phosphorylation sites on the Mcm2-7 tails promote DDK-dependent modulation of the number of CtGs formed per Mcm2-7. In a second, inefficient event, a subset of CtGs transfer their Cdc45 and GINS components to form CMGs. Importantly, higher CtG multiplicity results in increased frequency of CMG formation. Our findings reveal molecular mechanisms sensitizing helicase activation to DDK levels with implications for the control of replication origin efficiency and timing.

## Introduction

Eukaryotic DNA replication coordinates the assembly of multi-protein replisomes at origins of replication to ensure complete genome duplication. Replisome assembly begins during G1 phase when two copies of the Mcm2-7 helicase are loaded around origin DNA as a head-to-head double hexamer (Yardimci and Walter, 2014). In S phase, the association of helicase-activating factors selects a subset of these helicase double hexamers to initiate DNA unwinding and form the core of a bidirectional pair of replisomes (Mantiero et al., 2011, Tanaka et al., 2011). Correct temporal control of helicase activation both ensures that the genome is replicated exactly once per cell cycle (Remus and Diffley, 2009) and reserves a subset of origins to complete replication even if replisomes derived from adjacent origins stall (Blow et al., 2011; Mantiero et al., 2011).

Helicase activation is the committed step of replication initiation and is controlled during the cell cycle by two kinases: S-CDK and DDK (Figure 1 - figure supplement 1; Labib, 2010). DDK phosphorylates the long unstructured N-terminal tails of Mcm4 and Mcm6 driving the recruitment of Sld3/7 and Cdc45 to Mcm2-7 double hexamers (Kamimura et al., 2001; Kanemaki and Labib, 2006; Sheu and Stillman, 2010; Heller et al., 2011; Randell et al., 2010; Deegan et al., 2016). S-CDK phosphorylates Sld2 and Sld3/7 leading them to bind to two distinct pairs of BRCT repeats in Dpb11 (Tanaka et al., 2007; Zergerman and Diffley, 2007; Muramatsu et al., 2010). These S-CDK-dependent interactions recruit GINS to Sld3/7 associated with loaded Mcm2-7 complexes. Although both Cdc45 and GINS recruitment involve Sld3/7, whether these proteins are recruited by the same or different Sld3/7 molecules is unknown. Cdc45 and GINS are direct activators of the Mcm2-7 complex and association of these factors is required to form a pair of Cdc45-Mcm2-7-GINS complexes (CMGs), the active eukaryotic replicative helicase (Ilves et al., 2010; Moyer et al., 2006; Pacek et al., 2006). Finally, Mcm10 activates CMG complexes to unwind origin DNA (van Deursen et al., 2012; Kanke et al., 2012; Lõoke et al., 2017, Douglas et al., 2018). The result of this process is the formation of two oppositely-directed, active CMG complexes that generate the ssDNA necessary to recruit the rest of the replisome (Bell and Labib, 2016).

Although clear in broad strokes, multiple elements of CMG formation and helicase activation remain poorly understood. For example, current studies suggest that there are many Sld3-binding sites on the N-terminal tails of Mcm4 and Mcm6 although a specific binding motif has not been defined (Deegan et al., 2016). However, whether one or multiple sites are used during a given helicase-activation event is unknown. It is also unclear whether Cdc45 recruitment to the two Mcm2-7 complexes in the double hexamer is concerted. The possibility of coordinated Cdc45 recruitment is raised by evidence suggesting that Sld3 and Sld7 form a heterotetramer that could facilitate simultaneous recruitment of two Cdc45s (Itou et al., 2015). Lastly, the levels of DDK and a subset of helicase-activation proteins are known to modulate the timing and efficiency of helicase activation (Mantiero et al., 2011; Tanaka et al., 2011). However, the attributes of helicase activation that sensitizes this process to the levels of these proteins is not understood. To answer these questions and investigate the kinetics of the events leading to helicase activation, we developed a single-molecule (SM) method to directly observe this process in real time at a defined origin of replication *in vitro*.

Single-molecule biochemical techniques are well equipped to address questions of kinetics and stoichiometry during multi-protein complex assembly events like formation of the eukaryotic replisome (Stratmann and van Oijen, 2014). Previous single-molecule studies have complemented both structural and biochemical studies on helicase loading (Ticau et al., 2015; Ticau et al., 2017; Champasa et al., 2019). Like most complex multi-protein assembly processes, ensemble helicase-activation reactions occur asynchronously due to the large number of protein-DNA and protein-protein interactions involved. In addition, these bulk assays are typically performed as end-point assays with limited kinetic information (Yeeles et al., 2015), preventing detection of transient events. In contrast, single-molecule biochemical studies allow post-hoc synchronization of asynchronous events permitting detailed kinetic analysis. In addition, by allowing real-time visualization, single-molecule approaches can detect short-lived protein-protein or protein-DNA interactions, and determine precise relative stoichiometries.

Here we describe single-molecule biochemical assays for CMG formation and helicase activation that recapitulate previously demonstrated dependence on Sld3/7 and DDK. Initial recruitment of Cdc45 to the Mcm2-7 double hexamer occurs in a stepwise fashion. We observe many Cdc45 and GINS binding to individual Mcm2-7 double hexamers. Importantly, DDK levels modulate the number of these binding events and the frequency of final CMG formation. Consistent with these initial binding events being intermediates formed on the DDK-phosphorylated Mcm2-7 N-terminal tails, deleting the Mcm6 N-terminal tail reduces the number of these intermediates and the efficiency of CMG formation. Our findings support a model in which helicase activation is controlled by a combination of DDK-dependent regulation of an initial complex between Cdc45, GINS and the Mcm2-7 tails (the CtG complex) and inefficient conversion of the CtG to CMG complexes.

## RESULTS

### A single-molecule assay for CMG formation and DNA unwinding

We used Colocalization Single-Molecule Spectroscopy (CoSMoS, Friedman et al., 2006; Hoskins et al., 2011) to observe the process of eukaryotic CMG formation. This SM approach monitors the colocalization of fluorescently-labeled proteins with individual surface-tethered fluorescently-labeled DNA molecules to measure protein-DNA binding events. Fluorescent labeling of Mcm2-7, Cdc45, and GINS was accomplished using a SNAP-tag (Mcm2-7, Gendreizig et al., 2003, Ticau et al., 2015) or sortase-mediated coupling of fluorescent peptides (GINS, Cdc45, and Mcm2-7; Popp et al., 2007; Ticau et al., 2015, see Methods). These fluorescent tags did not interfere with protein function, as assessed by ensemble CMG-formation assays (Figure 1 - figure supplement 2). To prevent loaded Mcm2-7 complexes from sliding off the template prior to CMG formation (Ervin et al., 2009; Remus and Diffley, 2009), we used a 1.2 kb circular DNA containing a single origin of replication. Comparison of the circular template with a 1.3 kb linear template (Ticau et al., 2015) showed that the circular template exhibited a two-fold higher occupancy by Mcm2-7^4SNAP549^ at the end of a 20 min SM helicase-loading assay (Figure 1 - figure supplement 3).

To perform the CMG-formation reaction in a SM format, we executed a series of sequential incubations and monitored protein fluorescence (Figure 1A). After determining the locations of surface-coupled DNA molecules, we sequentially loaded the Mcm2-7^4SNAP549^ helicase, phosphorylated the loaded helicases with DDK, and then added the remaining proteins necessary to form the CMG (Figure 1 - figure supplement 1, S-CDK, Sld3/7, Cdc45, Dpb11, Sld2, GINS, Pol ε; Yeeles et al., 2015; Heller et al., 2011; Kanemaki and Labib, 2006; Kamimura et al., 2001). To ensure that only loaded Mcm2-7 molecules were present during CMG formation, we included a high-salt wash (HSW1) after DDK phosphorylation to remove helicase-loading intermediates. We initially focused on association of the Cdc45 component of CMG by labeling this protein with a distinct fluorophore (Cdc45^SORT649^). To focus our observations on the events of CMG formation after helicase loading, we only monitored protein fluorescence after DDK phosphorylation of loaded Mcm2-7^4SNAP549^ double hexamers. During this period, each DNA molecule was continuously monitored for fluorescent-protein colocalization for ∼30 min. After the CMG-formation reaction, we performed a second high-salt wash (HSW2) to distinguish fully-formed, salt-resistant CMG complexes from intermediates that are readily removed by this treatment (Yeeles et al., 2015).

**Figure 1.**
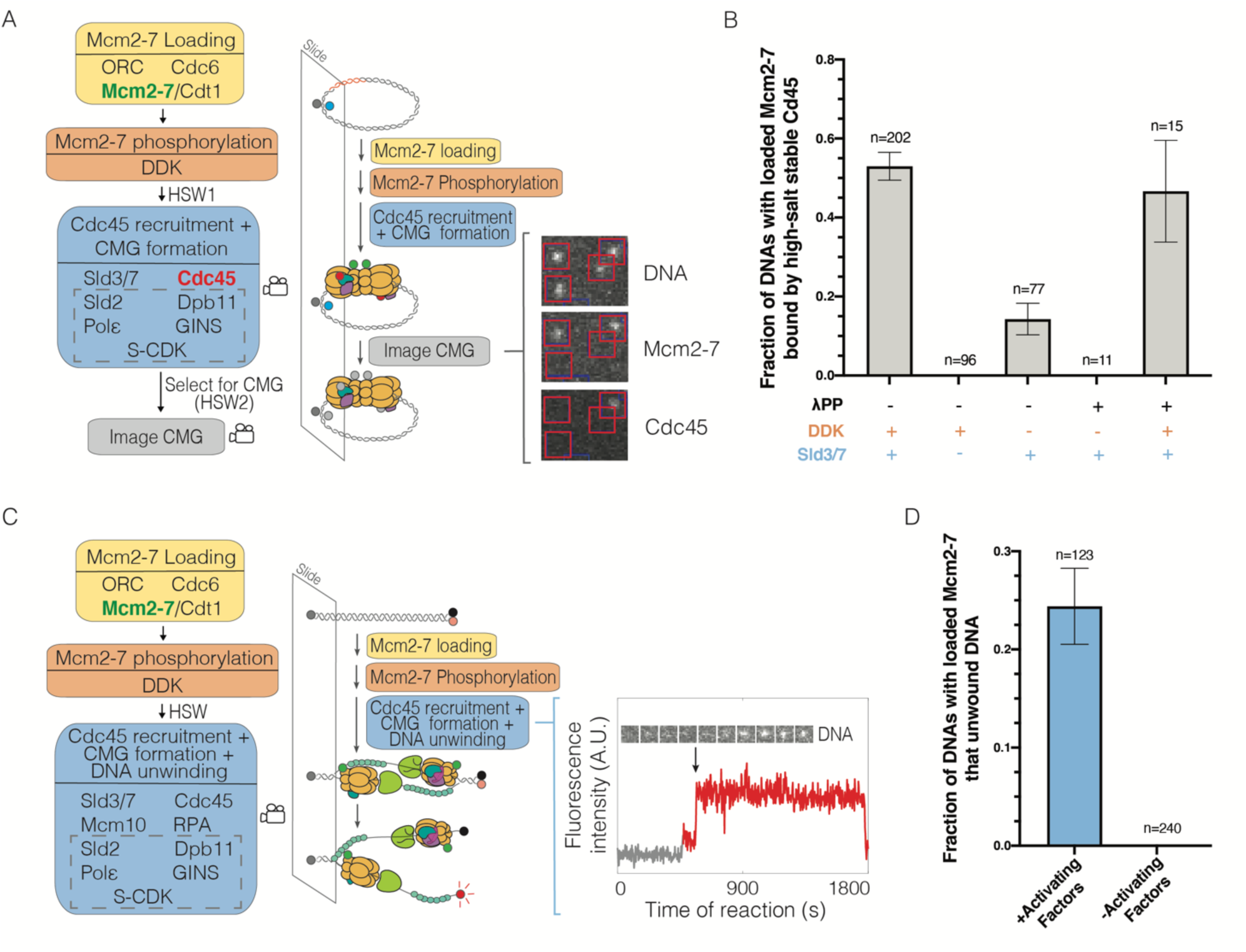
Single-molecule reaction for CMG formation and DNA unwinding. **(A)** Schematic for the single-molecule CMG formation reaction. Alexa-Fluor-488-labeled (blue circle) circular origin DNA molecules were tethered to the slide surface. Purified Mcm2-7^4SNAP549^ (green circle), Cdc45^SORT649^ (red circle), and other indicated proteins necessary to form the CMG were incubated with slide-coupled DNA in the steps shown. Members of the group of proteins referred to as SDPGC are in the dashed box. A high-salt wash (HSW1) was performed after Mcm2-7 phosphorylation to remove helicase-loading intermediates. Colocalization of fluorescently-labeled proteins with fluorescently-labeled DNA was monitored (indicated by camera icon) during CMG formation and after a second high-salt wash (HSW2). Example images show a small subregion of the microscope field of view taken at a single time point after the CMG formation reaction recorded in the color channels for DNA^488^, Mcm2-7^4SNAP549^, and Cdc45^SORT649^. Red squares are centered on DNA locations. Spots within the red square indicates stable binding of Mcm2-7^4SNAP549^ or Cdc45^SORT649^ after HSW2. **(B)** Cdc45 binding depends on Sld3/7 and DDK phosphorylation. The fraction (± SE) of n DNA molecules with bound Mcm2-7^4SNAP549^ that exhibited high-salt resistant Cdc45^SORT649^ at the first time point after the HSW2 is plotted for each of the conditions. DNAs are counted if they contain Mcm2-7^4SNAP549^ at the beginning of the CMG formation reaction. Where indicated, lambda phosphatase (λPP) was used to dephosphorylate Mcm2-7^4SNAP549^ prior to the Mcm2-7 loading reaction. **(C)** Schematic of the single-molecule DNA-unwinding reaction. Cy5- and BHQ-2-labeled (red and black circles) linear origin DNA molecules were coupled to microscope slides. The same stepwise incubations as with the single-molecule CMG-formation assay were used except Mcm10 and RPA were added to the “CMG formation and DNA unwinding step”. The plot displays a representative fluorescence intensity record for DNA^Cy5^ (increase in fluorescence indicated by arrow). An objective image-analysis algorithm (Friedman and Gelles, 2015) detected a spot of DNA fluorescence at time points shown in red. **(D)** DNA unwinding is dependent on activating factors. Helicase-activating factors eliminated were: S-CDK, Sld2, Dpb11, Pol ε, GINS, Sld3/7, Cdc45, Mcm10, and RPA. The fraction (± SE) of n DNA molecules that were unwound is plotted for both conditions.

Analysis of the DNA-associated proteins after the experiment showed multiple hallmarks of CMG formation. Consistent with efficient CMG formation, more than half (0.53 ± 0.04) of the DNAs with loaded Mcm2-7^4SNAP549^ complexes were bound with Cdc45^SORT649^ when observed immediately after the HSW2 high-salt wash (Figure 1B). These binding events were eliminated in the absence of Sld3/7, consistent with Cdc45^SORT649^ recruitment to Mcm2-7 relying on Sld3/7 (Figure 1B). Similarly, when Cdc6 was eliminated from the first step of the reaction to prevent helicase loading, subsequent CMG formation was dramatically reduced (fraction of DNAs bound by Cdc45 was 0.12 ± 0.01 versus 0.006 ± 0.006, Figure 1 - figure supplement 4). Thus, CMG formation was dependent on prior helicase loading. In the absence of DDK, high-salt-resistant bound Cdc45 was reduced but not eliminated (Figure 1B). We hypothesized that these “DDK-independent” binding events were due to residual DDK phosphorylation present on Mcm2-7 complexes purified from G1-arrested cells (Heller et al., 2011; Tanaka et al., 2011). Consistent with this idea, phosphatase treatment of Mcm2-7^4SNAP549^ prior to helicase loading eliminated CMG formation in the absence of DDK (Figure 1B). We confirmed that phosphatase-treated Mcm2-7^4SNAP549^ remained functional by sequential treatments with phosphatase and DDK. This sequence of treatments rescued CMG formation both in the SM (Figure 1B) and ensemble (Figure 1 - figure supplement 5) settings, demonstrating that the observed CMG formation was fully DDK-dependent.

To assess the functionality of the CMGs assembled in the SM setting, we asked whether they were able to unwind DNA. To this end, we generated a linear origin-containing DNA template with a fluorescent dye and a non-fluorescent quencher (BHQ-2) attached to opposing DNA strands at the surface-distal end of the DNA (Kose et al., 2019; Figure 1C). DNA unwinding promotes separation of the fluorescent dye-attached strand from the quencher-attached strand leading to increased dye emission. We simultaneously monitored labeled Mcm2-7^4SNAP549^ and unquenching of the DNA-associated fluorophore in the presence of the proteins required for CMG formation and two additional proteins (Mcm10 and RPA) required for extensive DNA unwinding (Douglas et al., 2018). Consistent with previous *in vitro* assays that showed only a fraction of loaded Mcm2-7 are activated (Douglas et al., 2018), we observed increased fluorescence for a fraction of DNAs with bound Mcm2-7^4SNAP549^ (0.24 ± 0.04, Figure 1D). Importantly, elimination of helicase-activating factors (S-CDK, Cdc45, GINS, Dpb11, Pol ε, Sld3/7, RPA, and Mcm10) from the reaction prevented all unquenching events (Figure 1D). Together, these studies demonstrate that our SM assays for CMG formation and helicase activation exhibit the outcomes and protein dependencies expected based on previous *in vivo* and ensemble *in vitro* studies (Kanemaki and Labib, 2006; Ilves et al., 2010; Heller et al., 2011; Tanaka et al., 2011; Yeeles et al., 2015).

### Cdc45 binding events to loaded Mcm2-7 are sequential

We next investigated the protein requirements for recruitment of Cdc45^SORT649^ to the Mcm2-7^4SNAP549^-DNA complex. Initially, we tested a minimal set of proteins predicted to recruit Cdc45 to loaded Mcm2-7 (Kanemaki and Labib, 2006; Heller, 2011; Tanaka et al., 2011). In these experiments, after phosphorylation of loaded Mcm2-7^4SNAP549^ with 1.3 nM DDK, we added only Sld3/7 and Cdc45^SORT649^ (Figure 2A). Under these conditions, we detected dynamic binding of one or two Cdc45^SORT649^ with the majority of Mcm2-7^4SNAP549^-bound DNA molecules (136/198, Figure 2A and Figure 2 - figure supplement 1). A subset (62/198) of DNAs with loaded Mcm2-7^4SNAP549^ never showed bound Cdc45. Consistent with the dynamic nature of the observed Cdc45 interactions and the requirement for additional proteins for CMG formation, in a separate experiment we did not detect high-salt-resistant Cdc45 binding to Mcm2-7 (0/70).

**Figure 2.**
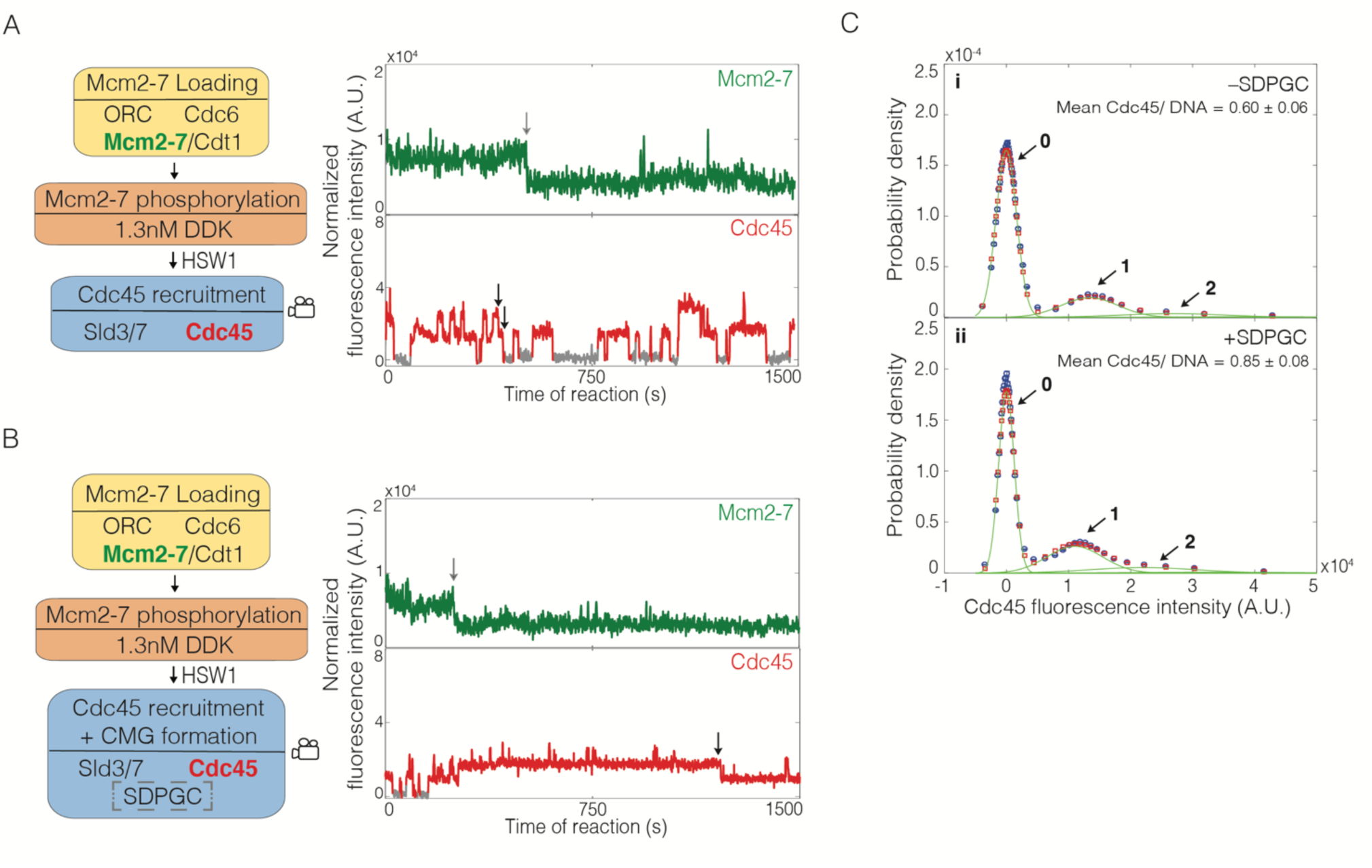
Sld2, Dpb11, Pol ε, GINS, and S-CDK (SDPGC) increases the frequency of Cdc45 binding events to the Mcm2-7 double hexamer. **(A)** Experiment protocol and representative fluorescence-intensity record for Mcm2-7^4SNAP549^ and Cdc45^SORT649^ at an origin-DNA location with minimal set of proteins required to recruit Cdc45. An objective image-analysis algorithm (Friedman and Gelles, 2015) detected a spot of protein fluorescence at time points shown in red or green. Photobleaching of Mcm2-7^4SNAP549^ is marked by a gray arrow and a set of likely dissociations of Cdc45^SORT649^ molecules are marked by black arrows. **(B)** Experiment protocol and representative fluorescence-intensity record for Mcm2-7^4SNAP549^ and Cdc45^SORT649^ at an origin-DNA location with all the factors required for CMG formation (+SDPGC = Sld2, Dpb11, Pol ε, GINS, and S-CDK). Photobleaching of Mcm2-7^4SNAP549^ is marked by a gray arrow and a potential dissociation of Cdc45^SORT649^ molecules is marked by a black arrow. **(C)** Multiple Cdc45^SORT649^ molecules bind to Mcm2-7^4SNAP549^-bound DNA. Cdc45^SORT649^ fluorescence**-**intensity histograms (blue) are shown for two conditions: –SDPGC (i) and + SDPGC (ii). The histogram data were fit to a sum-of-Gaussians model (red; see Materials and Methods). Fit parameters and calculated area fractions of individual Gaussian components (green) corresponding to the presence of the indicated numbers of Cdc45^SORT649^ molecules are given in Supplementary file 1 - table 1. The mean (± SE) numbers of Cdc45 molecules per DNA calculated from the fit parameters and the fraction Cdc45 labeled are indicated.

To assess the number of Cdc45 molecules bound to loaded Mcm2-7 more accurately, we constructed histograms that compiled Cdc45^SORT649^ intensities measured at all Mcm2-7^4SNAP549^-bound DNAs in every frame of the recorded images. We modeled these histogram data as the sum of equally spaced Gaussian peaks representing molecules with zero, one, two, or more bound Cdc45^SORT649^. The model assumed independent binding of Cdc45 to multiple identical sites on Mcm2-7 (see Materials and Methods). Based on this modeling, we determined the fraction of time a specific number of Cdc45^SORT649^ proteins were bound to DNAs with loaded Mcm2-7^4SNAP549^ (Figure 2A and Figure 2C, panel i). This analysis showed that one Cdc45^SORT649^ was bound to a DNA with an Mcm2-7^4SNAP549^ double hexamer 0.222 ± 0.002 of the time and two Cdc45^SORT649^ were bound only 0.079 ± 0.001 of the time (Supplementary file 1 - table 1). We also used this analysis to determine the average number of Cdc45 proteins bound to Mcm2-7^4SNAP549^. Taking into consideration our finding that 0.73 ± 0.07 of the Cdc45 protein was labeled (see Materials and Methods), we found that, on average, less than one Cdc45 molecule (labeled plus unlabeled) was associated with a loaded Mcm2-7 double hexamer (0.60 ± 0.06, Figure 2C, panel i).

We considered the possibility that the stepwise increases in Cdc45^SORT649^ fluorescence intensity (e.g., Figure 2A) represented transitions between states with two and four Cdc45 molecules instead of states with one and two molecules. If this were the case, however, we would expect to see a large subset of the bound Cdc45 to show half the fluorescent intensity relative to other events on the same DNA molecule. The reason for this expectation is that only a 0.73 ± 0.07 fraction of the Cdc45 protein is fluorescently labeled. Given this percentage of protein labeling, if the changes in fluorescent intensities were consistently due to simultaneous binding of two Cdc45 molecules, we would expect one of those molecules to be unlabeled 2*P* (1 – *P*) = 0.40 of the time, where *P* is the fraction labeled. Given the multiplicity of Cdc45 binding events observed on most DNAs, such a situation would result in two levels of increases in fluorescent intensity in any given trace. Instead, we observed Cdc45^SORT649^ fluorescence intensity changes that were consistently similar across a given trace (Figure 2A, bottom panel and Figure 2 - figure supplement 1). There are some events that might represent dimers of Cdc45 binding (e.g., Figure 2A, bottom, ∼900 s), but these are rare and are likely to be caused by two individual Cdc45^SORT649^ molecules binding in rapid succession. Thus, we conclude that the multiple Cdc45s binding to Mcm2-7-bound DNA are recruited as monomers in a sequential fashion.

We then assessed how Cdc45^SORT649^ binding events changed with the addition of the **S**ld2, **D**pb11, **P**ol ε, **G**INS, and S-**C**DK (SDPGC) proteins, all of which are required for CMG formation (Figure 2B and Figure 2 - figure supplement 2). We added these proteins to an otherwise unchanged reaction using 1.3 nM DDK. Fluorescent analysis showed that the addition of the SDPGC proteins lowered the fraction of time that DNA molecules bound by Mcm2-7^4SNAP549^ were not associated with Cdc45^SORT649^ (–SDPGC 0.682 ± 0.001; +SDPGC, 0.560 ± 0.001, Supplementary file 1 - table 1, rows 1 and 2). Consistent with this change, the average number of Cdc45 proteins (labeled plus unlabeled) on a Mcm2-7^4SNAP549^-bound DNA increased in these conditions (–SDPGC 0.60 ± 0.06; +SDPGC, 0.85 ± 0.08, Figure 2C, panel i and ii). The addition of the SDPGC proteins did not alter the uniformity of Cdc45^SORT649^ fluorescence intensity increases, consistent with Cdc45 still being recruited as a monomer in a sequential fashion (Figure 2B and Figure 2 - figure supplement 2). In contrast to SM experiments lacking the SDPGC proteins that showed no high-salt-resistant Cdc45 binding (0/70), the presence of the SDPGC proteins results in a fraction of loaded Mcm2-7 complexes forming the high-salt resistant Cdc45 complexes characteristic of CMG formation (0.15 ± 0.05, Figure 3A). Together, these results show that the complete set of proteins required for CMG formation leads to additional Cdc45 binding to loaded Mcm2-7, a subset of which go on to form CMG complexes.

**Figure 3.**
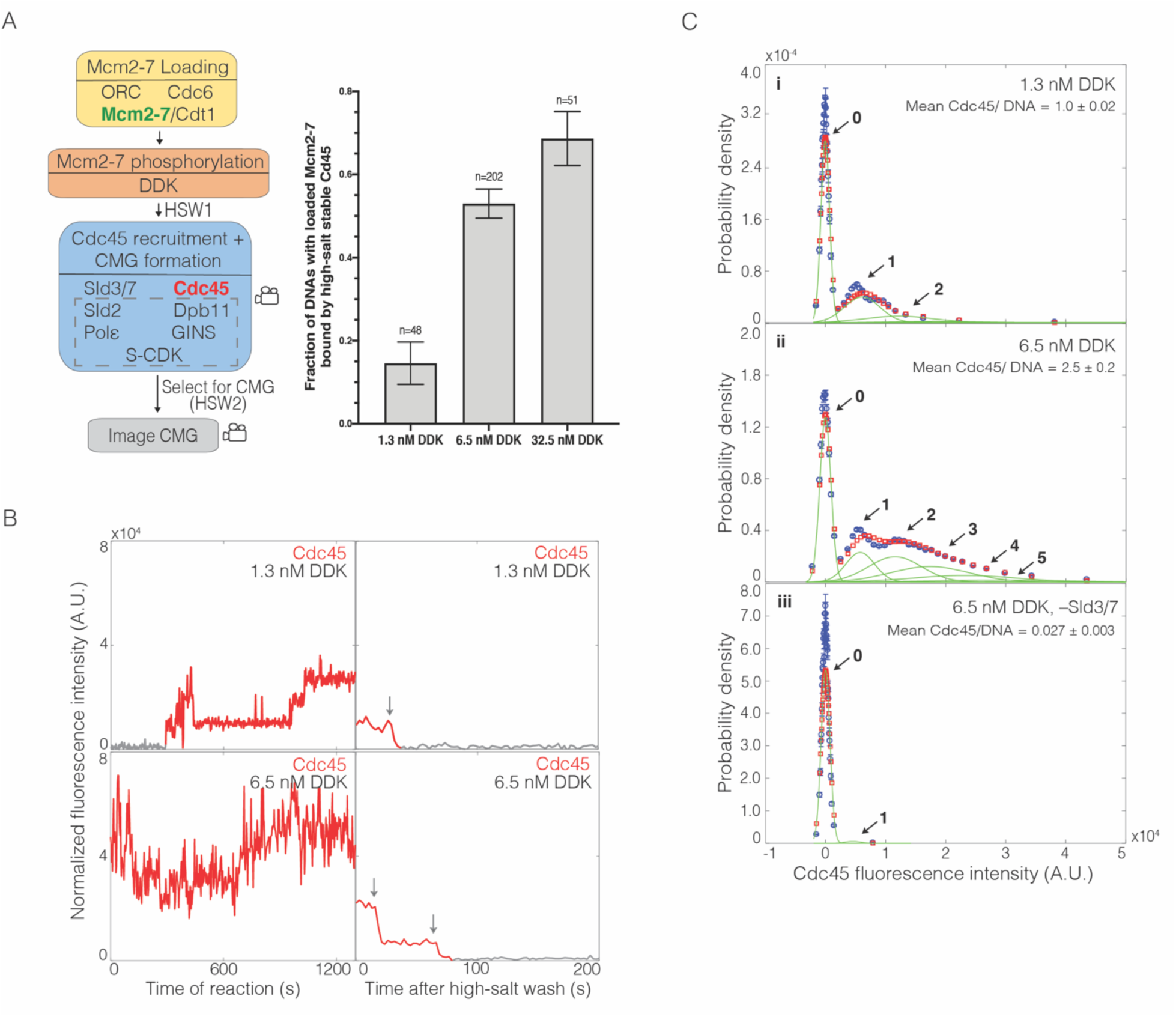
DDK modulates number of Cdc45 binding events and CMG formation efficiency. **(A)** Increasing DDK concentration results in more efficient CMG formation. The fraction of n (± SE) DNA molecules with loaded Mcm2-7^4SNAP549^ that were bound by high-salt stable Cdc45^SORT649^ are reported for reactions using 1.3 nM, 6.5 nM, or 32.5 nM DDK. Experimental protocol was as in Figure 1A. The data from 6.5 nM DDK condition is the same as Figure 1B, left bar. **(B)** Representative traces for Cdc45^SORT649^ under 1.3 nM and 6.5 nM DDK concentrations. Left panel is during CMG formation and right panel is after high-salt wash (HSW2). Points colored as in Figure 2B. Photobleaching of Cdc45^SORT649^ after the HSW2 is marked by gray arrows. Fluorescence intensities after HSW2 were adjusted to account for the higher excitation laser intensity used during photobleaching. **(C)** Multiple Cdc45^SORT649^ molecules bind to Mcm2-7^4SNAP549^. Cdc45^SORT649^ fluorescence**-**intensity histograms (blue) are shown for three conditions: (i) 1.3 nM DDK; (ii) 6.5 nM DDK; and (iii) 6.5nM DDK in the absence of Sld3/7. The histogram data were fit to a sum-of-Gaussians model (red; see Materials and Methods). Fit parameters and calculated area fractions of individual Gaussian components (green) corresponding to the presence of the indicated numbers of Cdc45^SORT649^ molecules are given in Supplementary file 1 - table 1. The mean (± SE) numbers of Cdc45 molecules per DNA calculated from the fit parameters and the fraction Cdc45 labeled are indicated.

### DDK levels modulate the frequency of CMG formation

Our early experiments with all of the proteins required for CMG formation yielded a relatively low fraction of loaded Mcm2-7 complexes being converted to CMGs. These results led us to consider approaches to increase the efficiency of CMG formation. Because higher DDK levels improve origin initiation efficiency (Mantiero et al., 2011; Tanaka et al., 2011), we explored the impact of increasing DDK concentration. At the higher DDK concentration used in Figure 1 (five-fold increase from 1.3 nM to 6.5 nM), the fraction of Mcm2-7^4SNAP549^-bound DNAs that became bound by high-salt resistant Cdc45^SORT649^ increased from 0.15 ± 0.05 to 0.53 ± 0.04 (Figure 3A). Elevating DDK concentration five-fold further (to 32.5 nM), increased the fraction of loaded Mcm2-7 complexes forming CMGs from 0.53 ± 0.04 to 0.69 ± 0.06 (Figure 3A). Because the relative increase in CMG formation was more dramatic between 1.3 nM and 6.5 nM (suggesting we were approaching saturation at 32.5 nM), we focused our subsequent studies on these two DDK concentrations.

Increased DDK levels led to each Mcm2-7^4SNAP549^-bound DNA interacting with more Cdc45^SORT649^ proteins during the CMG formation reaction (Figure 3B, left panels). Fluorescence intensity analysis revealed that elevating DDK levels to 6.5 nM increased the average number of Cdc45 molecules (labeled plus unlabeled) bound to Mcm2-7 by more than two-fold (1.0 ± 0.1 vs. 2.5 ± 0.2; Figure 3C, compare panels i and ii). This analysis also revealed sometimes four or more Cdc45^SORT649^ molecules bound to an individual Mcm2-7^4SNAP549^ double hexamer at 6.5 nM DDK. Importantly, all of the Cdc45^SORT649^ binding events remain Sld3/7-dependent, suggesting that the Cdc45^SORT649^ binding detected requires indirect binding with the Mcm2-7 tails via Sld3/7 (Figure 3C, panel iii). The fraction of DNAs with Mcm2-7^4SNAP549^ bound at least once by Cdc45 is similar between the 1.3 nM (40 out of 48 molecules, 0.83 ± 0.05) and 6.5 nM (71/87, 0.82 ± 0.06) DDK conditions, indicating that increasing DDK is primarily impacting the number of Cdc45 molecules bound per double hexamer rather than the fraction of double hexamers that are capable of interacting with Cdc45. Finally, we note that although these data give a sense of the multiplicity of Cdc45 proteins bound to Mcm2-7 at a specific time, these binding events are dynamic. Thus, the number of Cdc45 binding events to an Mcm2-7 double hexamer over the course of the reaction is higher.

The multiplicity of the bound Cdc45 suggests that binding occurs on the Mcm4 and Mcm6 tails rather than the core of the Mcm2-7 complex. The core of each Mcm2-7 molecule is thought to have only one site that directly binds Cdc45 (the site of Cdc45 association in the CMG, Yuan et al., 2016, Goswami et al., 2018). In contrast, there are many Sld3 binding sites on both the Mcm4 and Mcm6 N-terminal tails (Deegan et al., 2016), each of which could recruit Cdc45. Thus, the appearance of more than two bound Cdc45 molecules indicates that at least a subset (and perhaps most, see below) of the Cdc45 binding events observed under high DDK conditions reflect Cdc45 binding (via Sld3) to the phosphorylated N-terminal tails.

### Only a subset of the multiple Mcm2-7-bound Cdc45 proteins form CMGs

To determine the number of Cdc45^SORT649^ proteins retained after the second high-salt wash (HSW2, Figure 1A), we increased the laser intensity and monitored protein fluorescence until all the protein-bound dyes were photobleached. Because photobleaching is stochastic, each photobleaching event indicates the presence of one fluorescently-labeled protein. Although our intensity model suggests that a given Mcm2-7 is capable of simultaneously binding four or more Cdc45 molecules during the CMG formation reaction (Figure 3C, panel ii), we never (zero of 202 examined) observed that more than two Cdc45^SORT649^ proteins remained bound to a Mcm2-7-bound DNA after the HSW2 high-salt wash (e.g., Figure 3B, right panel). Although it is possible in principle that we undercounted photobleaching events that were not temporally resolved, comparison of the rate of photobleaching under these conditions to the time resolution of the experiment indicated that this would be rare (Figure 3 - figure supplement 1). The maximum of two high-salt resistant Cdc45s is consistent with the interpretation that high-salt resistance indicates that the Cdc45s are incorporated into CMGs (Yeeles et al., 2015). Thus, although increased DDK leads to more Cdc45 proteins binding simultaneously to each Mcm2-7-bound DNA, at most two are incorporated into the final CMG complexes.

We frequently observed only one high-salt stable Cdc45^SORT649^ associated with Mcm2-7-bound DNA, leading us to ask whether CMG formation occurred independently or was coordinated for the two Mcm2-7 complexes present in a loaded double hexamer. If CMG formation was coordinated for the two Mcm2-7 complexes, we would expect most of the high-salt resistant Cdc45 to occur in pairs. Using the measured fraction labeled of Cdc45^SORT649^ (*P* = 0.73 ± 0.07), we calculated that if CMG formation was coordinated, 0.57 ± 0.09 (*P*^2^) of the bound high-salt resistant Cdc45^SORT649^ should include two labeled Cdc45s. In contrast, we observed that only 42/102 (0.41 ± 0.05) of the Mcm2-7-bound DNAs with salt-resistant Cdc45 included two labeled Cdc45s when CMG formation was performed with 6.5 nM DDK. When the same measurement is performed with 1.3 nM DDK, this fraction drops to 1/7 (0.14 ± 0.13), consistent with lower numbers of bound Cdc45 leading to more frequent formation of a single CMG within the Mcm2-7 double hexamer. These findings suggest that CMG formation on the two Mcm2-7 complexes in each double hexamer can occur independently.

### Cdc45 dwell times are similar at low and high DDK levels

We next asked whether the complex pattern of Cdc45 binding events observed at high-DDK levels represented a fundamentally different type of Cdc45 association, or an increased frequency of the same type of Cdc45 association observed at lower DDK levels. To distinguish between these two possibilities, we repeated the SM CMG-formation assay with Cdc45 having a reduced labeling fraction (0.036 ± 0.004, see Materials and Methods). These conditions allowed us to observe binding of individual labeled Cdc45 molecules even in the high-DDK context when many additional (unlabeled) Cdc45 were simultaneously bound to the same Mcm2-7-bound DNA (Figure 4A; Figure 4 - figure supplement 1). Having established conditions in which we could consistently observe single Cdc45 molecules bound to Mcm2-7-bound DNA, we determined the distributions of bound Cdc45 dwell times (Figure 4B). We did not see a significant difference in the dwell time distributions of Cdc45^SORT649^ at 1.3 nM and 6.5 nM DDK, consistent with the idea that high-DDK levels increases the number of Cdc45 associations to Mcm2-7-bound DNA without changing their properties.

**Figure 4.**
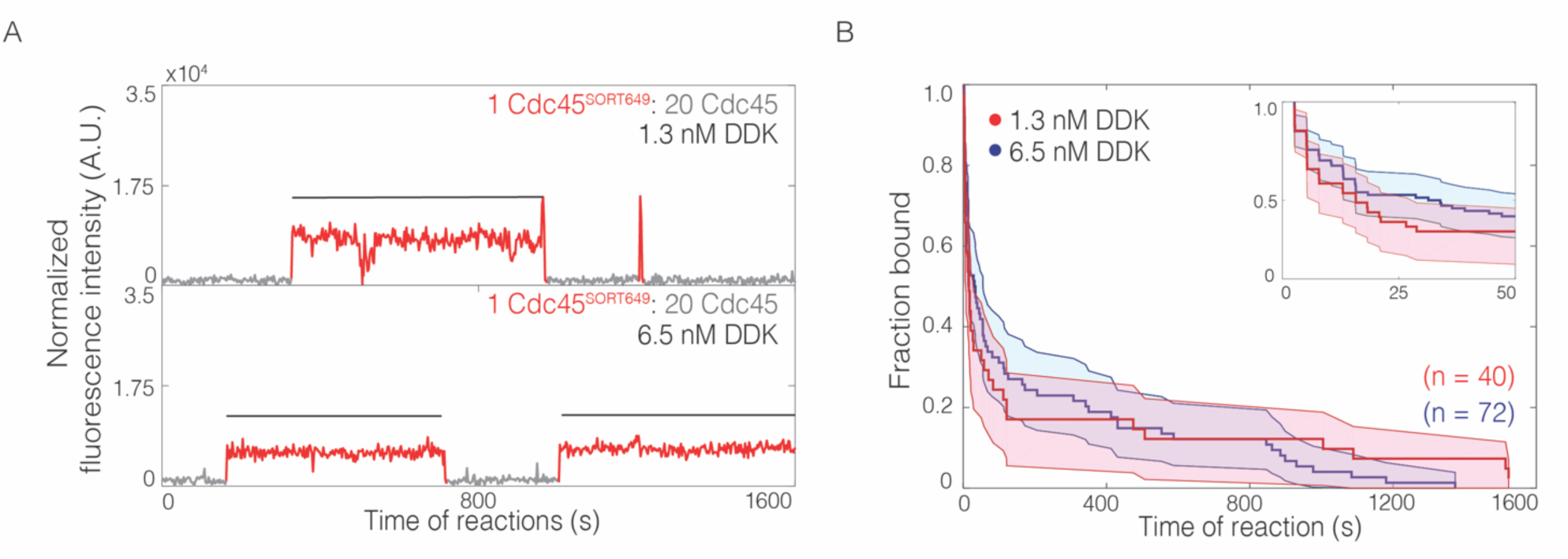
The lifetimes of individual bound Cdc45 molecules are not significantly changed by DDK levels. **(A)** Representative traces for Cdc45^SORT649^ under conditions in which only 0.036 ± 0.004 of Cdc45 molecules were fluorescently labeled at 1.3 nM (top) and 6.5 nM (bottom) DDK. Points colored as in Figure 2B. Black line indicates single Cdc45^SORT649^ binding. **(B)** Survival function for Cdc45^SORT649^ dwell times on Mcm2-7-bound DNA. The vertical axis represents the fraction of Cdc45^SORT649^ molecules that remain bound after the dwell interval indicated on the horizontal axis. Shaded areas represent the 95% confidence intervals for each curve. Inset: magnified view.

### DDK levels modulate the number of Mcm2-7 tail phosphorylation events

Elevating DDK levels could change Mcm2-7 phosphorylation in multiple ways. More DDK could increase the number of phosphates per Mcm2-7 complex as well as the fraction of Mcm2-7 complexes that have any phosphates. DDK is known to exhibit stable binding to Mcm2-7 (Sheu and Stillman, 2006; Ramer et al., 2013; Abd Wahab and Remus, 2020), and an extreme model predicts that when DDK binds to Mcm2-7 this interaction leads to complete Mcm4 and Mcm6 tail phosphorylation. If this was the case, then DDK concentration would only modulate the fraction of Mcm2-7 complexes being completely phosphorylated, not the number of phosphates per Mcm2-7 tail. To investigate these possibilities, we exposed loaded Mcm2-7 complexes to increasing concentrations of DDK and monitored the extent of phosphorylation of Mcm4 and Mcm6 using Phos-tag™ gels (Figure 4 - figure supplement 2). Changes in the mobility of the Mcm4 and Mcm6 indicated a distribution of numbers of phosphates for both proteins, arguing against the latter model in which only complete tail phosphorylation occurs. Instead, as DDK concentration increases we see evidence for both increasing numbers of phosphates on the Mcm4 and Mcm6 proteins and a higher fraction of the molecules being modified. These findings are consistent with DDK concentration regulating the number of Cdc45 binding through modulation of the number of Mcm4 and Mcm6 tail phosphorylation events.

### Cdc45-dependent binding of multiple GINS to Mcm2-7

To investigate the incorporation of GINS into CMGs, we performed a SM assay for CMG formation in which GINS was fluorescently labeled (Figure 5A). We first determined the number of GINS^SORT649^ retained after HSW2. Consistent with GINS incorporating into CMGs, we observed that the fraction of Mcm2-7-bound DNAs with bound, high-salt-resistant GINS^SORT649^ increased when the concentration of DDK was raised from 1.3 to 6.5 nM (0.19 ± 0.04 vs 0.72 ± 0.04; Figure 5A). As with Cdc45, we never observed more than two high-salt stable GINS^SORT649^ after HSW2 (Figure 5B), and it is unlikely that we missed additional binding events due to simultaneous photobleaching of two GINS^SORT649^ molecules (Figure 5 - figure supplement 1). In addition, elimination of Sld3/7 from the assay resulted in a 10-fold reduction in high-salt resistant GINS^SORT649^ bound to Mcm2-7^2SORT549^-bound DNA (0.07 ± 0.02; Figure 5A). We next investigated the binding of GINS to Mcm2-7 during CMG formation. Because GINS is thought to be recruited by indirect interactions (via Dpb11 and Sld2) with S-CDK-phosphorylated Sld3 (Tanaka et al., 2007; Zegerman and Diffley, 2007; Muramatsu et al., 2010), it was possible that we would observe the same DDK-phosphorylation-controlled multiplicity of GINS complexes binding to loaded Mcm2-7 that we observed for Cdc45. Alternatively, it was possible that GINS recruitment could require interactions with the structured regions of Mcm2-7 involved in CMG formation. In that case, we would not observe more than two GINS bound. Consistent with the first model, we observed dynamic binding of many GINS^SORT649^ to Mcm2-7^2SORT549^-bound DNA molecules (Figure 5B). As with Cdc45, the multiplicity of bound GINS^SORT649^ was strongly dependent on the presence of Sld3/7 (compare Figure 5C, panels ii and iii) and Mcm2-7 (Figure 5 - supplement 2).

**Figure 5.**
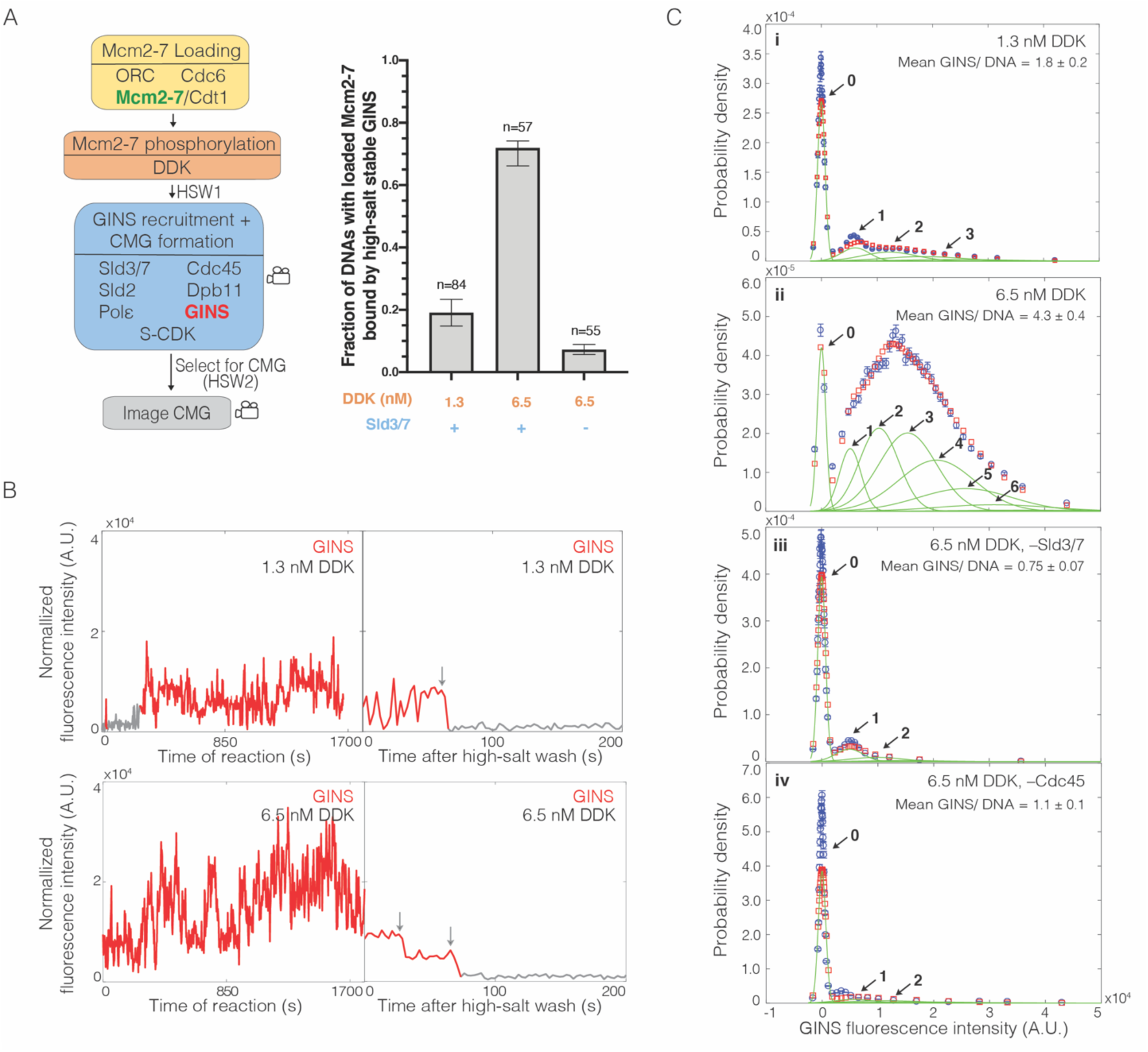
Multiple GINS bind to Mcm2-7. **(A)** Increasing DDK concentration results in more efficient CMG formation that is dependent on the presence of Sld3/7. Left: schematic representation of the experiment. Right: graph showing the fraction of n Mcm2-7^2SORT549^-bound DNA molecules that were bound by high-salt stable GINS^SORT649^ under the indicated conditions. **(B)** Representative GINS^SORT649^ fluorescence intensity records at two DNA molecules in the presence of 1.3 nM or 6.5 nM DDK. Left panel is during CMG formation and right panel is the same molecule after the second high-salt wash (HSW2). Points colored as in Figure 2B. Photobleaching of GINS^SORT649^ after HSW2 is marked by gray arrows. **(C)** GINS^SORT649^ fluorescence**-**intensity histograms (blue) are shown for four conditions: (i) 1.3 nM DDK; (ii) 6.5 nM DDK; (iii) 6.5 nM DDK in the absence of Sld3/7; and (iv) 6.5 nM DDK in the absence of Cdc45. The histogram data were fit to a sum-of-Gaussians model (red; see Materials and Methods). Fit parameters and calculated area fractions of individual Gaussian components (green) corresponding to the presence of the indicated numbers of GINS^SORT649^ molecules are given in Supplementary file 2 - table 1. The mean (± SE) numbers of GINS molecules per DNA calculated from the fit parameters and the fraction GINS labeled are indicated.

The patterns of GINS binding during CMG formation support the hypothesis that GINS is initially recruited to the Mcm4 and Mcm6 tails. Fluorescence intensity analysis revealed primarily two or fewer GINS^SORT649^ molecules binding to a loaded Mcm2-7^2SORT549^ double hexamer (Figure 5C, panel i) at 1.3 nM DDK. Taking into account the fraction labeled of GINS (0.71 ± 0.07), this analysis indicated an average of 1.8 ± 0.2 total GINS molecules (labeled plus unlabeled) bound per Mcm2-7-bound DNA (Figure 5C, panel i). In contrast, at 6.5 nM DDK, five or more GINS^SORT649^ complexes could bind to an Mcm2-7^2SORT549^-DNA complex with an average of 4.3 ± 0.4 GINS (labeled plus unlabeled) per Mcm2-7-bound DNA (Figure 5C, panel ii). This average number of bound GINS^SORT649^ is inconsistent with a model where all GINS interactions take place at the two GINS binding sites involved in CMG formation present in a double hexamer. As with Cdc45, the fraction of DNAs with Mcm2-7^2SORT549^ that bound GINS^SORT649^ at least once was similar for 1.3 nM (76/84, 0.90 ± 0.04) and 6.5 nM (54/57, 0.95 ± 0.03). Because almost all complexes bound GINS under both conditions, the impact of DDK on CMG formation is clearly due to increased numbers of GINS bound per Mcm2-7 double hexamer.

Current models suggest that GINS is recruited by a series of CDK-dependent protein binding events involving Sld3, Dpb11, and Sld2 (Bell and Labib, 2016, Figure 1 - figure supplement 1). This mechanism raises the possibility that different Sld3 molecules could recruit Cdc45 and GINS. To investigate this possibility, we performed the CMG formation assay with GINS^SORT649^ but in the absence of Cdc45. We observed dramatically lower numbers of bound GINS (Figure 5C, compare panels ii and iv), comparable to those observed in the absence of Sld3/7 (Figure 5C, compare panels iii and iv). Thus, we conclude that the initial binding of GINS requires both Sld3/7 and Cdc45 and that the recruitment of GINS and Cdc45 occurs via the same Sld3/7 molecules.

### Loss of Mcm6 N-terminal tail reduces CMG formation

Together, our data support a model in which complexes including both Cdc45 and GINS form at multiple sites on the DDK-phosphorylated Mcm2-7 N-terminal tails. We refer to these complexes as Cdc45-tail-GINS (CtG) complexes. Our data also suggests that increasing the numbers of CtGs formed increases the probability of CMG formation. Based on these observations, we propose that the conversion of CtGs into CMG complexes is inefficient with multiple CtGs releasing prior to successful CMG formation.

To test the model that the multiple associations of Cdc45 involve interactions with DDK-phosphorylated Mcm4 and Mcm6, we asked how deleting the Mcm6 N-terminal tail impacted this event. We focused on the Mcm6 N-terminal tail because deletion of the Mcm4 N-terminal tail is lethal (Champasa et al., 2019). We constructed a fluorescently-labeled Mcm2-7 complex that lacked the Mcm6 N-terminal tail (Mcm2-7^4SNAP549-6ΔN^). Using Mcm2-7^4SNAP549-6ΔN^ and Cdc45^SORT649^, we monitored the pattern of Cdc45 binding and the extent of CMG formation using 6.5 nM DDK in the SM assay.

Consistent with the Mcm6 N-terminal tail contributing to CtG formation, use of the Mcm2-7^4SNAP549-6ΔN^ mutant in our assay resulted in reduced Cdc45^SORT649^ binding even at 6.5 nM DDK (compare Figure 6A with Figure 3B, bottom panel). Compared to WT Mcm2-7^4SNAP549^ at either 1.3 or 6.5 nM DDK, the mutant at 6.5 nM DDK showed a lower fraction of loaded Mcm2-7 molecules that were converted to CMGs (Figure 6B). Thus, reducing the numbers of these initial recruitment events by eliminating the Mcm6 N-terminal tail clearly alters the efficiency of CMG formation. Consistent with the low number of CMGs, the average number of Cdc45 (labeled plus unlabeled bound to Mcm2-7^4SNAP549-6ΔN^ during CMG formation at 6.5 nM DDK was also low (0.10 ± 0.01, Figure 6C). Together, these data indicate that the Mcm6 N-terminal tail contributes to recruiting or retaining multiple Cdc45 proteins. Given the dependence of GINS association on Cdc45, we infer that binding of GINS is also dependent on the Mcm6 N-terminal tail. Thus, deletion of the Mcm6 N-terminal tail reduces both the initial recruitment of Cdc45 and the formation of CMGs.

**Figure 6.**
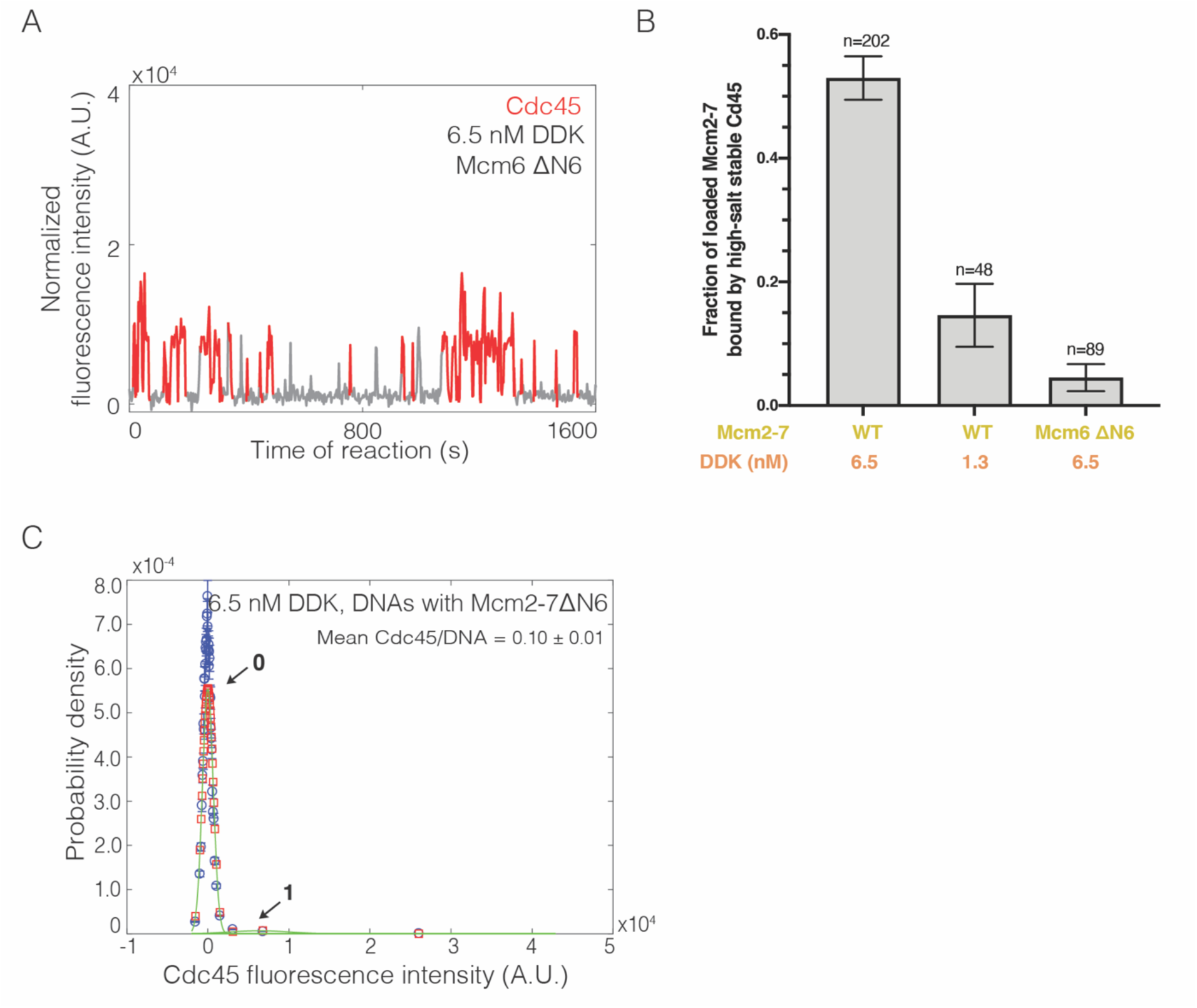
Loss of the Mcm6 N-terminal tail reduces Cdc45 binding events and CMG formation. **(A)** Representative trace for Cdc45^SORT649^ binding to Mcm2-7^4SNAP549-6ΔN^-bound DNA during CMG formation at 6.5 nM DDK. Points colored as in Figure 2B. **(B)** Deletion of the Mcm6 N-terminal tail reduced high-salt stable Cdc45 binding to Mcm2-7-bound DNAs, indicative of CMG formation. The data from 1.3 nM and 6.5 nM DDK are the same as Figure 3A. **(C)** Binding of Cdc45^SORT649^ to Mcm2-7^4SNAP5496ΔN^-bound DNA. Fit parameters and calculated area fractions of individual Gaussian components (green) corresponding to the presence of the indicated numbers of Cdc45^SORT649^ molecules are given in Supplementary file 1 - table 1. The mean (± SE) number of Cdc45 molecules per DNA calculated from the fit parameters and the fraction Cdc45 labeled are indicated.

## Discussion

### Cdc45 and GINS are recruited by the Mcm2-7 N-terminal tails

The studies presented here provide evidence for a new intermediate in the helicase-activation process, the CtG, which can form at multiple sites on the Mcm2-7 N-terminal tails. We found that DDK levels modulate the number of these intermediates formed per loaded Mcm2-7 complex and that only a subset of CtG transition to a CMG (Figure 7). We propose that the combination of these properties create a mechanism that sensitizes helicase activation to DDK activity, in order to control replication initiation efficiency.

**Figure 7.**
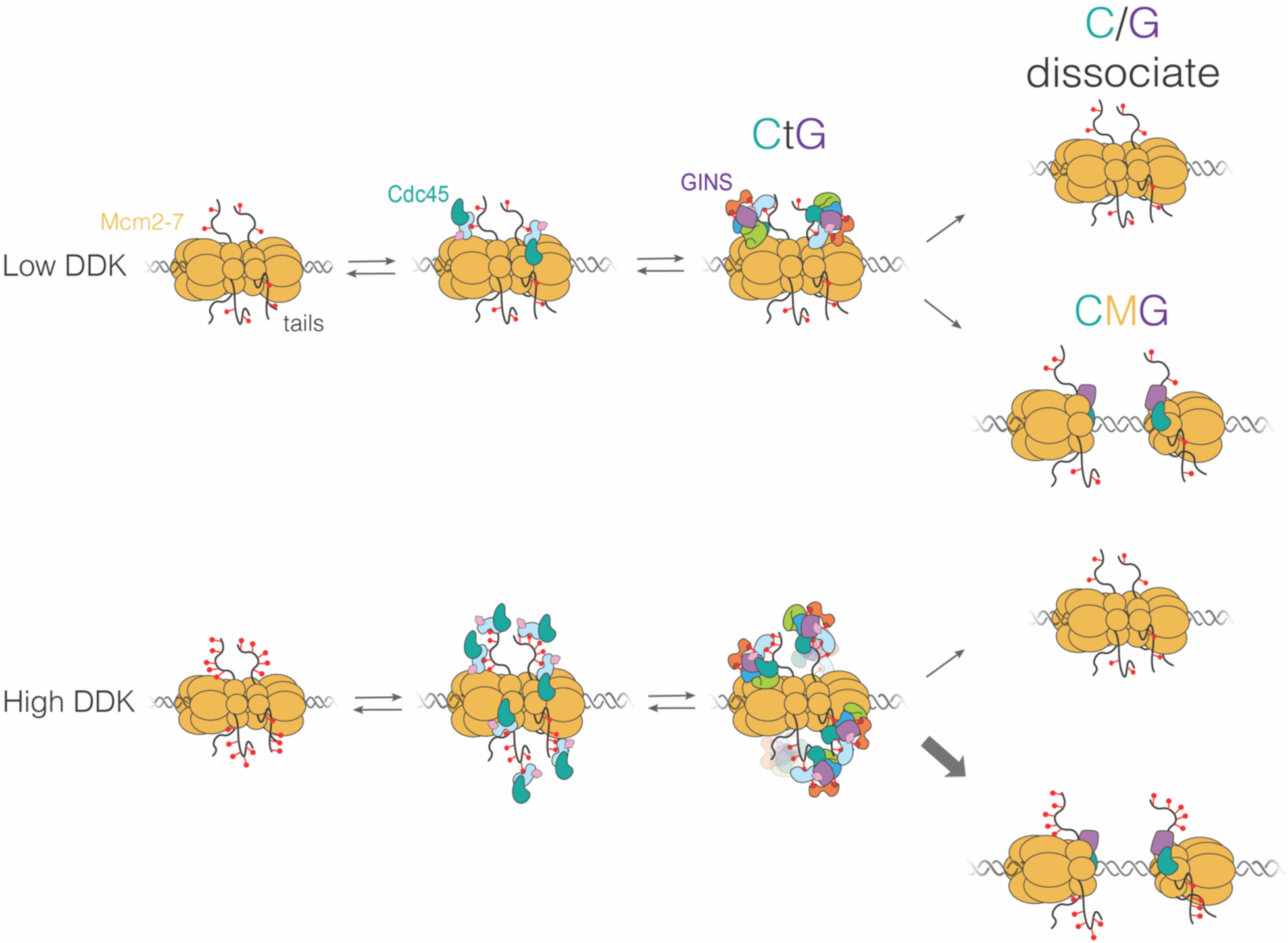
Proposed model for CMG formation. DDK levels control the amount of Mcm4 and Mcm6 N-terminal tail phosphorylation. These modifications indirectly (via Sld3/7) recruit Cdc45 and GINS, forming the CtG intermediate. The CtG intermediate can then follow one of two paths: a) dissociation, or b) deposition onto the structured core of the Mcm2-7 helicase to form the CMG. We hypothesize that the rate of this conversion is the same for any CtG, but it is infrequent. Thus, having more CtGs (e.g., at higher DDK levels) increases the probability of CMG formation, which we assume is irreversible. Relevant proteins are labeled in the illustration; red lollipops represent phosphorylations; CtG = Cdc45-tails-GINS; CMG = Cdc45-Mcm2-7-GINS; C/G = Cdc45-GINS.

Previous studies found that Sld3 contains a phosphopeptide-binding domain that recognizes both DDK-dependent and DDK-independent phosphorylation sites on Mcm4 and Mcm6 N-terminal tails (Deegan et al., 2016). Binding of Sld3 to the DDK-dependent phosphorylation sites is required for the recruitment of Cdc45 to the Mcm2-7 double hexamer (Deegan et al., 2016). It was not clear, however, if these interactions involved only the Mcm4 and Mcm6 tails or additional sites on Mcm2-7, how they led to subsequent CMG formation, or how the many potential binding sites for Sld3/7 on Mcm4 and Mcm6 tails contributed.

Here we present several lines of evidence that support the initial recruitment of both GINS and Cdc45 involve formation of a multi-protein complex with the Mcm2-7 N-terminal tails. First, there are only two non-tail binding sites on Mcm2-7 for Cdc45 or GINS, and yet we see more than two of these proteins bound during CMG formation (Figure 3 and Figure 5). Second, because the Mcm4 and Mcm6 N-terminal tails are the targets of DDK and Sld3 binding (Sheu and Stillman, 2006; Randell et al., 2010; Deegan et al., 2013), the dependence of the multiple Cdc45 and GINS binding events that we observe on DDK activity and Sld3/7 is consistent with a Mcm2-7 tail binding site (Figure 1B, Figure 3C, and Figure 5). Third, the dwell time distributions of Cdc45 associations at high or low DDK levels are similar (Figure 4), arguing that the increased multiplicity of Cdc45 binding is not due to a distinct mechanism of Cdc45 association at high DDK levels. Fourth, eliminating the Mcm6 N-terminal tail, greatly reduces CtG formation (Figure 6). Finally, the strong dependence of GINS recruitment on the presence of Cdc45 is consistent with a simple mechanism in which these proteins are recruited sequentially to the tails to form a single complex rather than binding with the tails at independent positions (Figure 5C).

Our evidence also supports the CtG being an intermediate on the way to CMG formation. Consistent with this hypothesis, elimination of factors that promote CtG formation (e.g., DDK and Sld3/7) also inhibit CMG formation (Figure 1B and Figure 5A). Similarly, elimination of a subset of CtG binding sites by deleting the Mcm6 N-terminal tail, results in a strong reduction in CMG formation (Figure 6B). Indeed, throughout our studies we observe a strong correlation between the number of CtGs formed on a Mcm2-7 double hexamer and the efficiency of CMG formation.

### What are the components of the CtG?

Although our findings indicate that the CtG minimally includes Cdc45 and GINS bound to Mcm6 (and likely Mcm4) N-terminal tail(s), this complex is likely to contain other proteins involved in CMG formation. The robust Sld3/7 dependence of Cdc45 and GINS binding to Mcm2-7 (Figure 3 and Figure 5) suggests that the CtG includes this protein. Current models propose that a network of interactions between Sld3, Dpb11, Sld2, and GINS are required to form the CMG (Tanaka et al., 2007; Zegerman and Diffley, 2007; Muramatsu et al., 2010). Given Sld3’s known binding to phosphorylated N-terminal tails (Deegan et al., 2016), the simplest model is that these interacting proteins are required to form CtGs, and subsequently CMGs. It is possible that Pol ε is simultaneously recruited with GINS to the CtG as part of a preformed pre-LC complex (Muramatsu et al., 2010).

The full set of roles performed by Sld3, Dpb11, Sld2, and Pol ε in CtG and CMG formation remains to be determined. These proteins could be involved only in CtG assembly or their release could facilitate the transition of Cdc45 and GINS binding with the Mcm2-7 tails to form the final interactions characteristic of the CMG. Finally, the requirement of Cdc45 to recruit GINS (Figure 5C, panel iii) strongly suggests that the CtG involves direct interactions between these proteins, suggesting the interesting possibility that key interactions between these proteins involved in the CMG are already present in the CtG.

### DDK effects are mediated by inefficient CtG to CMG conversion

Our studies indicate that the conversion from CtG to CMG is inefficient. With low DDK levels, we often observed two Cdc45 proteins that associated with Mcm2-7 double hexamers over long periods of time (>100 sec; 33 long-lived binding events/48 double hexamers). However, only seven of these 33 converted to CMGs (Figure 3A), indicating that most of the CtGs either dissociate or fail to become part of the CMG before the end of the experiment (Figure 7, top). We hypothesize that this inefficient conversion is true at all DDK concentrations. When DDK concentration was raised, we observed increased multiplicity of CtG formation (Figure 3A and 5A). In contrast, increased DDK is not associated with a significant change in dwell times of individual Cdc45 molecules (Figure 4) or the fraction of Mcm2-7 double hexamers bound by Cdc45 or GINS. Based on these findings we propose that, due to the inefficient conversion of CtGs to CMGs, the increased multiplicity of CtGs on a given double hexamer at higher DDK levels leads to increased opportunities for CtG conversion to CMG (Figure 7, bottom) and, therefore, increased rates of origin activation.

### Implications for the control of cellular origin activation

Our model is consistent with studies addressing the control of origin initiation efficiency and replication timing (Mantiero et al., 2011; Tanaka et al., 2011). These studies showed that increasing the expression of a subset of limiting helicase-activation proteins, including the Dbf4 regulatory subunit of DDK, increased origin initiation efficiency and advanced the time of replication initiation for late initiating origins. Our findings suggest that these changes would increase the number of CtGs formed per double hexamer and consequently the efficiency of CMG formation. We propose that the combination of multiple potential phosphorylation sites on the Mcm4 and Mcm6 tails, and the inefficient conversion of the CtGs formed on the tails to the CMG, tunes the helicase-activation process to respond to levels of DDK and other helicase-activation factors. Consistent with this model, reducing the number of phosphorylation sites by deleting the Mcm6 tail reduced CtG and CMG formation at the same DDK levels (Figure 6).

Our proposed mechanism is analogous to the regulation of Sic1 proteolysis by CDK (Nash et al., 2001). Multiple CDK phosphorylation sites on Sic1 are each weakly recognized by the Cdc4 F-box protein, leading to ubiquitinylation and degradation of Sic1. Although weakly bound, the accumulation of multiple phosphorylation sites on Sic1 promotes a switch-like response to CDK levels. We propose that the similar mechanisms are operating during helicase activation, with the Mcm4 and Mcm6 tails providing the multiple phosphorylation-dependent binding sites and the inefficient conversion of the CtG to the CMG performing the role analogous to the suboptimal Cdc4 interaction.

Processes beyond those shown in Figure 7 are likely to play important roles in origin activation. For example, our experiments did not include phosphatases that could counteract the action of DDK and CDK. Such phosphatases might be recruited by Rif1, a protein that interacts with protein phosphatase 1 and has been implicated in the regulation of replication initiation (Hafner et al., 2018; Hiraga et al., 2014, 2018; Mattarocci et al., 2018). The presence of phosphatases could regulate Mcm2-7 N-terminal tail phosphorylation levels, further sensitizing origin activation to changes in DDK level. In addition, phosphatases may be essential to remove Mcm2-7-tail phosphorylation after helicase activation to ensure that the tails of active helicases do not continue to compete for limiting helicase-activation factors.

## Acknowledgements

We are grateful to Alexandra Pike and Shalini Gupta for comments on the manuscript, Karim Labib for gifting the plasmid pFJD5, and Michael Maloney for making the Cdc45 labeling construct.

## Author contributions

L.D.J.K. performed all experiments with feedback from J.G., L.J.F., and S.P.B. C.R. made constructs and partially purified mutant protein for Figure 6. L.D.J.K., L.J.F., and J.G. analyzed the data. S.P.B. and L.D.J.K. composed the paper with input from all authors. S.P.B. and J.G. directed the project.

## Materials and Methods

### Preparation of unlabeled proteins

Wild-type unlabeled Mcm2-7/Cdt1 and ORC complexes were purified as described previously (Kang et al., 2014). Wild-type Cdc6 was purified as described in Frigola et al. (2013). Wild-type DDK, S-CDK, Sld3/7, Sld2, unlabeled Cdc45, Dpb11, unlabeled GINS, Mcm10, Pol ε and RPA were purified as described in Lõoke et al. (2017).

### Preparation of GINS^SORT649^

Plasmid pFJD5 (gift from K. Labib) was modified with a His-Tag followed by a PreScission Protease tag at the N-terminus of Psf3 and a LPTEGG at the C-terminus of Psf3 (pLD01, Supplementary file 3–table 2) to facilitate peptide addition by Sortase. The plasmid was expressed in BL21 (D3) competent *E. coli* (NEB) and purified as described in Yeeles et al. (2015) with the following modifications. After eluting from HisPur Ni-NTA resin (ThermoFisher Scientific), the eluates were incubated with excess PreScission Protease at 4°C overnight, then incubated with HisPur Ni-NTA resin to remove any uncleaved GINS. The unbound protein was collected and applied to MonoQ column as described in Yeeles et al. (2015). The peak GINS fractions were collected and Sortase was used to attach the peptide NH_2_-GGGHHHHHHHHHHC-COOH coupled to maleimide-Dy649P1 (Dyomics) to the C-terminus of Psf3 to form GINS^SORT649^. Sortase coupling of the fluorescent peptide was performed as described in Ticau et al. (2017). After fluorescent labeling, GINS ^SORT649^ was applied to a Superdex S200 column as described in Yeeles et al. (2015). Peak fractions were pooled and applied to Ni-NTA resin pre-equilibrated with buffer H (25 mM HEPES-KOH [pH 7.6], 5 mM MgOAc, 10% glycerol, and 0.02% NP-40), supplemented with 200 mM potassium acetate (KOAc), 0.02% Nonidet P-40 (NP-40) and 10 mM imidazole, for 30 min with rotation at 4°C to separate peptide-coupled GINS from uncoupled GINS. The resin was washed with buffer H, supplemented with 200 mM KOAc, 0.02% NP-40 and 30 mM imidazole. Fluorescently labeled GINS^SORT649^ was eluted using buffer H, supplemented with 200 mM KOAc, 0.02% NP-40 and 250 mM imidazole. Peak fractions were pooled, aliquoted, and stored at −80°C.

### Preparation of Cdc45^SORT649^

A CDC45 expressing plasmid was modified by appending sequences encoding a Flag-epitope followed by LPETGG at the 5’ end of the gene of plasmid pMM032 (Supplementary file 3–table 2). This plasmid was integrated into genome of yMH109 to create yMM034 (Supplementary file 3–table 1). This strain was used to purify Cdc45 as described in Lõoke et al. (2017) with the following modifications. After elution from anti-Flag M2 affinity gel (Sigma), the eluate was attached with Sortase to the peptide NH_2_-GGGHHHHHHHHHHC-COOH coupled to maleimide-Dy649P1 (Dyomics) as described in Ticau et al. (2017). The modified protein was applied to a Superdex S75 column equilibrated with buffer H, supplemented with 0.3 M KGlut and 5% glycerol. The peak fractions were applied to a HisPur Ni-NTA resin (ThermoFisher Scientific) equilibrated with buffer H, supplemented with 300 mM KGlut and 5 mM imidazole, for 30 min with rotation at 4°C to separate peptide-coupled from uncoupled Cdc45. The flow-through was discarded and the resin was washed with buffer H, supplemented with 300 mM KGlut, and 20 mM imidazole. Fluorescently labeled Cdc45^SORT649^ was eluted using buffer H, supplemented with 300 mM KGlut and 250 mM imidazole. Peak fractions were pooled, aliquoted, and stored at −80°C.

### Preparation of Mcm2-7^4SNAP549^, Mcm2-7^4SNAP5496ΔN^, and Mcm2-7^2SORT549^

The Mcm6-Mcm7 overexpression plasmid was modified by deleting the sequences encoding Mcm6 amino acids 2 to 105 to create the plasmid pCKR006 (Supplementary file 3–table 2). This plasmid was integrated into the genome of a strain that contained expression constructs for Mcm2, Mcm3, Mcm4, Mcm5 and Cdt1 to create the yeast strain yCKR006 (Supplementary file 3–table 1). Mcm2-7/Cdt1 ΔN6 (yCKR006), Mcm2-7/Cdt1 SORT-tag (LPETGG at the C-terminus of Mcm2 for fluorescent labeling, yST179, Supplementary file 3–table 1), and Mcm2-7/Cdt1 SNAP-tag (SNAP-tagged at Mcm4 for fluorescent labeling, yST147, Supplementary file 3–table 1) were purified from the indicated yeast strains as described previously (Kang et al., 2014) with the following modifications. After ultracentrifugation, the whole cell extract was applied to 1 mL of anti-Flag M2 affinity gel (Sigma) pre-equilibrated with buffer H, 200 mM KGlut, 0.01% NP-40 and 1 mM ATP and incubated with rotation for 2 hrs at 4°C. The flow-through was discarded and the resin was washed with 30 mL of buffer H, supplemented with 300 mM KGlut, 0.01% NP-40 and 1 mM ATP. Mcm2-7/Cdt1 was eluted with buffer H, supplemented with 300 mM KGlut, 0.01% NP-40, 1 mM ATP and 0.1 mg/mL 3xFLAG peptide for the first elution and with 0.3 mg/mL 3xFLAG peptide for the consecutive elutions. After elution from anti-Flag M2 affinity gel, the eluate was fluorescently labeled as follows. For Mcm2-7^4SNAP549^ and Mcm2-7^4SNAP5496ΔN^ (SNAP at the N-terminus of Mcm4), the FLAG eluate was labeled with SNAP-Surface549 (NEB) by incubating 3x molar excess of dye at 4°C overnight. After coupling the protein to the fluorophore, the reaction was applied to a Superdex S200 column equilibrated with buffer H, supplemented with 300 mM KGlut, 0.01% NP-40, and 1 mM ATP. Peak fractions containing Mcm2-7^4SNAP5496 or 4SNAP5496ΔN^ /Cdt1 were pooled, aliquoted, and stored at −80°C. Mcm2-7^4SNAP549^/Cdt1 was used in all experiments with Cdc45^SORT649^, with exception of experiments with N-terminal mutant Mcm2-7.

For Mcm2-7^2SORT549^/Cdt1 (LPETGG at the C-terminus of Mcm2) the FLAG eluate was attached with Sortase to the peptide NH_2_-GGGHHHHHHHHHHC-COOH coupled to maleimide-Dy649P1 (Dyomics) as described in Ticau et al. (2017). Mcm2-7^2SORT549^/Cdt1 was applied to a Superdex S200 column equilibrated with buffer H, supplemented with 300 mM KGlut, 0.01% NP-40, and 1 mM ATP. Peak fractions containing fluorescently labeled Mcm2-7^2SORT549^/Cdt1 were pooled, aliquoted, and stored at −80°C. Mcm2-7^2SORT549^/Cdt1 was used in all experiments with _GINS_SORT649.

### Determining Mcm2-7, Cdc45 and GINS labeling fraction

To determine what fraction of Mcm2-7^4SNAP549^ were fluorescently labeled, SNAP-Surface549 labeled Mcm2-7 was mixed with maleimide-DY-649P1 dissolved in anhydrous DMSO in a 1:1 ratio, and the reaction and analysis was carried out as described in Ticau et al. (2015). The labeled fraction for Mcm2-7^4SNAP549^ was determined to be 0.73 ± 0.07. To determine the labeling fraction of Cdc45^SORT649^ (and GINS^SORT649^) was mixed with maleimide-DY549P1 dissolved in anhydrous DMSO in a 1:1 molar ratio at 4°C for 10 min. The maleimide-DY549P1 would label all free cysteines in the protein. The reaction was terminated with 2 mM DTT. We added 5 nM of maleimide-DY549P1-labeled Cdc45^SORT649^ (or 8nM maleimide-DY549P1-labeled GINS^SORT649^) to the SM CMG-formation assay with 1.3 nM DDK to obtain single Cdc45^SORT649^ (or GINS^SORT649^) traces. The fraction of maleimide-DY549P1-labeled Cdc45^SORT649^ (or GINS^SORT649^) that also contained DY-649P1 was determined and reported as the percent labeling by the DY-649P1 (we assume that coupling of maleimide-DY-549P1 or DY-649P1 to Cdc45, Mcm2-7, and GINS are not influenced by the presence or absence of the 649 or 549 label). The labeled fraction for Cdc45^SORT649^ and GINS^SORT649^ were determined to be 0.73 ± 0.07 and 0.71 ± 0.07, respectively. To lower the labeling fraction of Cdc45^SORT649^ to 0.036, the Cdc45^SORT649^ preparation was mixed at a 1:19 ratio with unlabeled Cdc45.

### Generation of circular template

To create biotin- and fluorescently-labeled circular origin DNA, a 1.3 kb ARS1-containing interval of the DNA plasmid template pUC19-ARS1 was amplified with NotI-site containing primers, one of which contained a biotin and the other an Alexa-Fluor-488 dye. After PCR cleanup with QIAquick PCR Purification Kit (Qiagen), the PCR product was digested with NotI at 37°C for 4 hrs and repurified with a QIAquick PCR Purification Kit. We next performed a ligation with 0.2 ng/uL of digested DNA and 0.04 U/uL of T4 DNA ligase (NEB) at 18°C overnight to favor intramolecular ligation. The ligation products were purified by phenol-chloroform extraction and concentrated by ethanol precipitation. The concentrated DNA was then run on a 1.5% TBE-agarose gel and the circular DNA band was extracted and purified with QIAquick Gel Extraction Kit (Qiagen).

### Single-molecule assay for CMG formation

Biotinylated Alexa-Fluor-488-labeled, 1.2-kb-long circular-DNA molecules containing an origin were coupled to the surface of a reaction chamber through streptavidin. We identified DNA molecule locations by acquiring 4-7 images with 488 nm excitation at the beginning of the experiment. Helicase loading reaction was performed as described in Ticau et al. (2015). Reaction buffers for helicase loading were as described in Ticau et al. (2015). All subsequent steps (i.e., Mcm2-7 phosphorylation and CMG formation) used the same helicase-loading buffer. After helicase loading either 1.3 or 6.5 nM DDK was added. Chambers were then washed with two chamber volumes of buffer A (25 mM HEPES-KOH [pH 7.6], 5 mM magnesium acetate (MgOAc), 0.02% NP-40), supplemented with 0.5 M NaCl (HSW1), followed by one chamber volume of buffer A supplemented with 300 mM KGlut. Next, CMG-formation reaction was added containing 7.5 nM CDK, 15.5 nM Sld2, 10 nM Dpb11, 7.5 nM Pol ε, 50 nM GINS, 12.5 nM Sld3/7 and 12.5 nM Cdc45. DNA was imaged immediately after adding the CMG-formation reaction to the slide but not throughout the experiment. When using Cdc45^SORT649^ or GINS^SORT649^, they were added at 5 nM and 7 nM, respectively. After ∼30 min, chambers were washed with four chamber volumes of buffer A supplemented with 0.5 M NaCl (HSW2) followed by one chamber volume of buffer A supplemented with 300 mM KGlut.

### Single-molecule assay for DNA unwinding

To obtain the 1.1 kb linear-DNA molecules used for the DNA unwinding assay, the same origin-containing interval of pUC19-ARS1 was amplified with a NotI-site containing primer and a biotin-containing primer. The PCR-amplified DNA was digested with Not1, purified with a QIAquick PCR Purification Kit (Qiagen), and ligated with oligos each containing a Cy5- or BHQ-2 label and NotI sticky ends (NEB) complimentary to the amplified DNA template. After ligation, the DNA product was purified with a QIAquick PCR Purification Kit (Qiagen). The DNA was coupled to the surface of a reaction chamber in a streptavidin-mediated reaction to slide-attached PEG-biotin as described previously (Ticau et al, 2015).

Helicase-loading, DDK, and CMG-formation reactions were performed as described above with the following modifications: During CMG formation 12.5 nM RPA and 0.625 nM Mcm10 were added. No high-salt wash was performed after CMG-formation reaction.

### Single-Molecule Microscopy

The micro-mirror total internal reflection (TIR) microscope used for multiwavelength single-molecule using excitation wavelengths 488, 532, and 633 nm as described in Friedman et al., (2006). Briefly, the chamber surface was cleaned and derivatized using a 200:1 ratio of silane-NHS-PEG and silane-NHS-PEG-biotin as described in Ticau et al. (2015). Biotinylated AlexaFluor488-labeled 1.2kb circular DNA molecules containing an origin were coupled to the surface of a reaction chamber through streptavidin. Subsequently, Mcm2-7 loading, Mcm2-7 phosphorylation, and CMG-formation reactions (or DNA unwinding) were added to a chamber of the microscope slide. DNA was imaged immediately after adding the CMG-formation reaction to the slide but not throughout the experiment. After addition of CMG-formation proteins to Mcm2-7-bound DNA, frames of 1 sec duration were acquired according to the following protocol: (1) 1 sec frame with the 532 nm laser on and 1 sec frame with the 633 nm laser on, and (2) a computer-controlled focus-adjustment using a 785-nm laser (Crawford et al., 2008). This cycle was repeated over the course of a ∼30 min experiment. After ∼30 min of incubation, chambers were washed with four chamber volumes of buffer H supplemented with 500 mM NaCl and one chamber volume of buffer H supplemented with 300 mM of KGlut. Recording of protein photobleaching was then performed as described in Ticau et al. (2015).

### Single-molecule data analysis

Analysis of the CoSMoS data sets was as described in Ticau et al. (2015). Records of protein single-molecule fluorescence were corrected for background fluorescence as follows: One set of intensity time records summed intensities of 5 × 5 pixel areas centered on the DNA molecule locations (*D_j_*(*m*) records for integrated intensity from the *m*th frame for the *j*th DNA molecule area). Additional time records similarly recorded time-smoothed background intensity levels from areas adjacent to each DNA molecule location (*B_j_*(*m*) background records for the *j*th DNA molecule) and the time-smoothed background offset record for the camera with no light incident (*C*(*m*) record). Background-corrected fluorescence intensity records were calculated as *D_j_*(*m*) − *B_j_*(*m*).

### Intensity histogram fitting

Background fluorescence (solution dye + slide autofluorescence) results in a (*B_j_*(*m*) − *C*(*m*)) value that is proportional to the local laser excitation at the *j*th DNA site. One DNA site record was chosen as having a reference level of laser excitation (*B*_R_(*m*) − *C*(*m*)), and the time record for all other DNA sites were background-corrected and normalized to correspond to the laser excitation at that reference *j*=R site. Accordingly, this background-corrected normalized time record *N_j_*(*m*) for each DNA site is

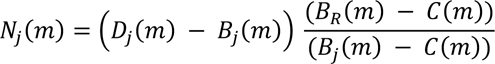

Using a noise (variance) vs. (signal mean) measurement we determined the calibration factor *k*, the number of photons corresponding to one digitized intensity unit (analog-to-digital unit, ADU) (*k* = 0.0118 photons/ADU) (Friedman et al., 2006). From the recording of each experiment, we defined *g = <*(*B*_R_(*m*) − *C*(*m*))> *k*, the mean value (in photons) of the reference DNA site background intensity averaged over all frames for that experiment.

The intensity data *N_j_*(*m*) from *j* = 200 to 400 DNA locations over *m* = 600 time points (1500 s recording duration) were pooled for intensity distribution analysis of *N*. Outlier measurements with *N* values > 50,000 ADU (in all experiments less than 5% of measurements) were excluded from histograms and fits.

The distributions of corrected intensities *N* from each experiment were fit to a modified Binomial-distributed Gaussian mixture model probability density function

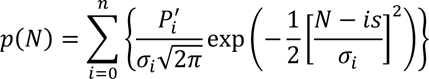

with the normalized amplitudes

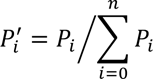

derived from the modified Binomial distribution

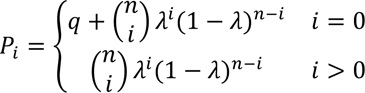

and the component widths

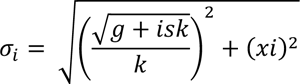

with fit parameters *q*, *s*, λ, and *x.* The number of components was held fixed at *n* = 8 although component numbers for *i* > 6 did not contribute significantly to the probability density function (2^$^ < 0.01) in any of the data sets analyzed (Supplementary file 1-table 1 and Supplementary file 2 - table 1). In this model *q* is an additional amplitude in the zeroth peak to account for target molecules incapable of binding labeled protein; *s* is the corrected intensity (expressed in ADU) corresponding to a single dye molecule; *n*λ is the mean number of dye molecules per binding-competent Mcm2-7-bound DNA; and *x* is additional variation in fluorescent spot intensity (expressed in ADU) presumed to arise from DNA Brownian motion. Fit parameters were determined by maximizing the likelihood using customized Matlab software (in https://github.com/gelles-brandeis/jganalyze).

### Bulk CMG-formation assay

Ensemble CMG-formation assays were performed as described in Champasa et al. (2019).

### Phosphorylation assays on SuperSep™ Phos-tag™ gels

Each incubation step was performed in a thermomixer (Eppendorf) with shaking at 1250 rpm at 25 °C. The DNA template was the same used in CMG-formation assays as described in Champasa et al. (2019). Helicase loading reactions contained 100 nM Mcm2-7, 45 nM ORC and 45 nM Cdc6. After loading, DDK was added at varying concentrations (0 nM, 20 nM, 80 nM, 120 nM, 260 nM and 500 nM) for 20 min. The supernatant was removed by applying the reaction to a DynaMag-2 magnet (ThermoFisher Scientific). Reactions were washed with buffer H, 300 mM KCl, and 0.01% NP-40 three times. Proteins were released from the DNA by incubation with 5 U of DNA I (Worthington) in 10 mL of 25 mM HEPES-KOH (pH 7.6), 5 mM MgOAc, 200 mM NaCl, 5% glycerol, 0.02% NP-40, and 2 mM CaCl_2_ for 20 min at 25 °C before running on a pre-cast 10 % SuperSep™ Phos-tag™ gel at 25 mA for >2hrs. The gels were then stained using Krypton™ Protein Stain (ThermoFisher Scientific).

When Mcm4 tail-SNAP protein was used, then 500 nM of protein was phosphorylated with varying concentrations of DDK (0 nM, 20 nM, 80 nM, 120 nM, 260 nM and 500 nM) for 20 min. Reactions were run on a pre-cast 10 % SuperSep™ Phos-tag™ gel at 25 mA for >2hrs. The gels were then stained using Krypton™ Protein Stain (ThermoFisher Scientific).

## Supplementary information

**Figure 1–figure supplement 1.**
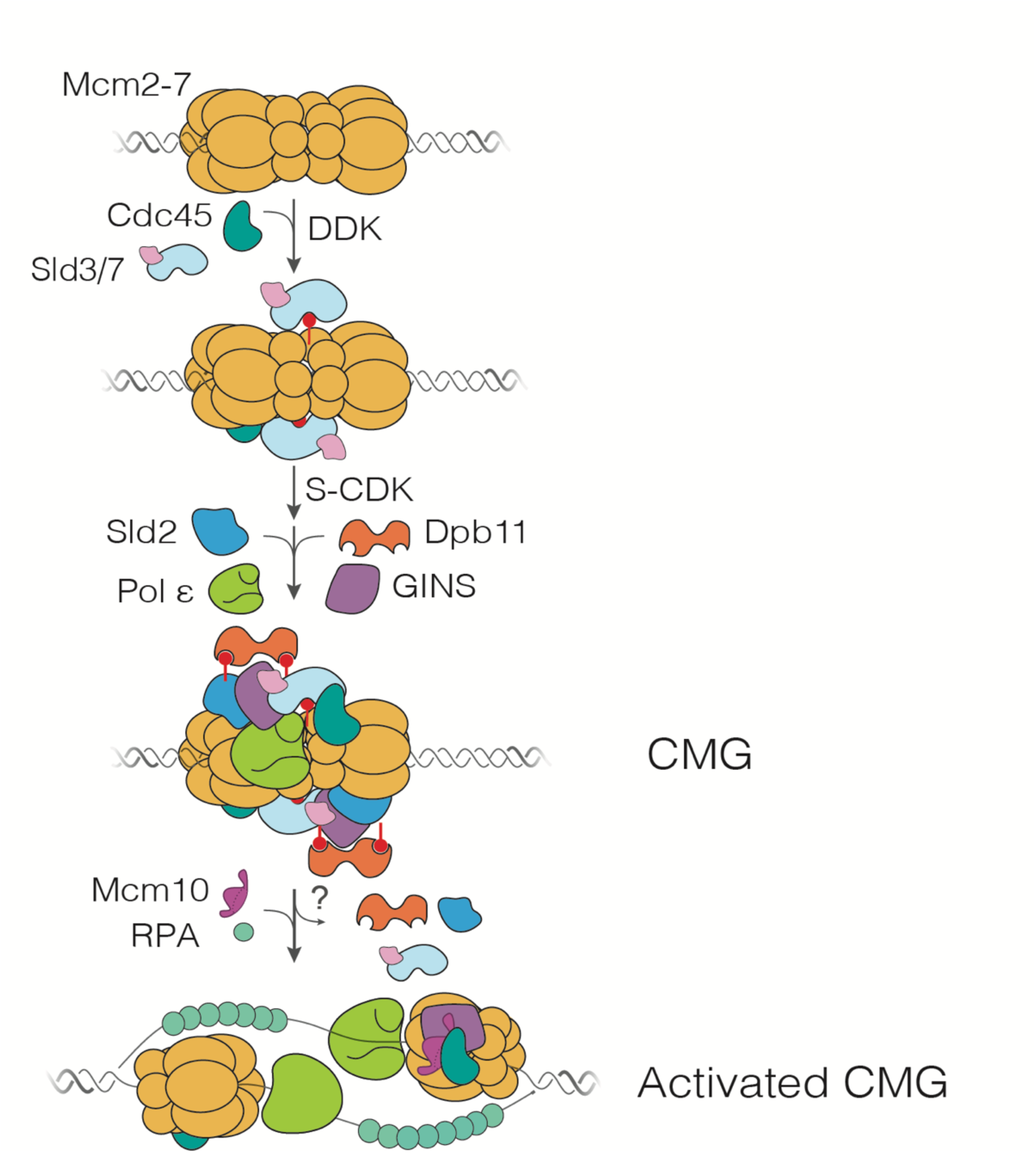
Schematic representation of DNA replication initiation. Scheme shows the first time that each factor is required.

**Figure 1–figure supplement 2.**
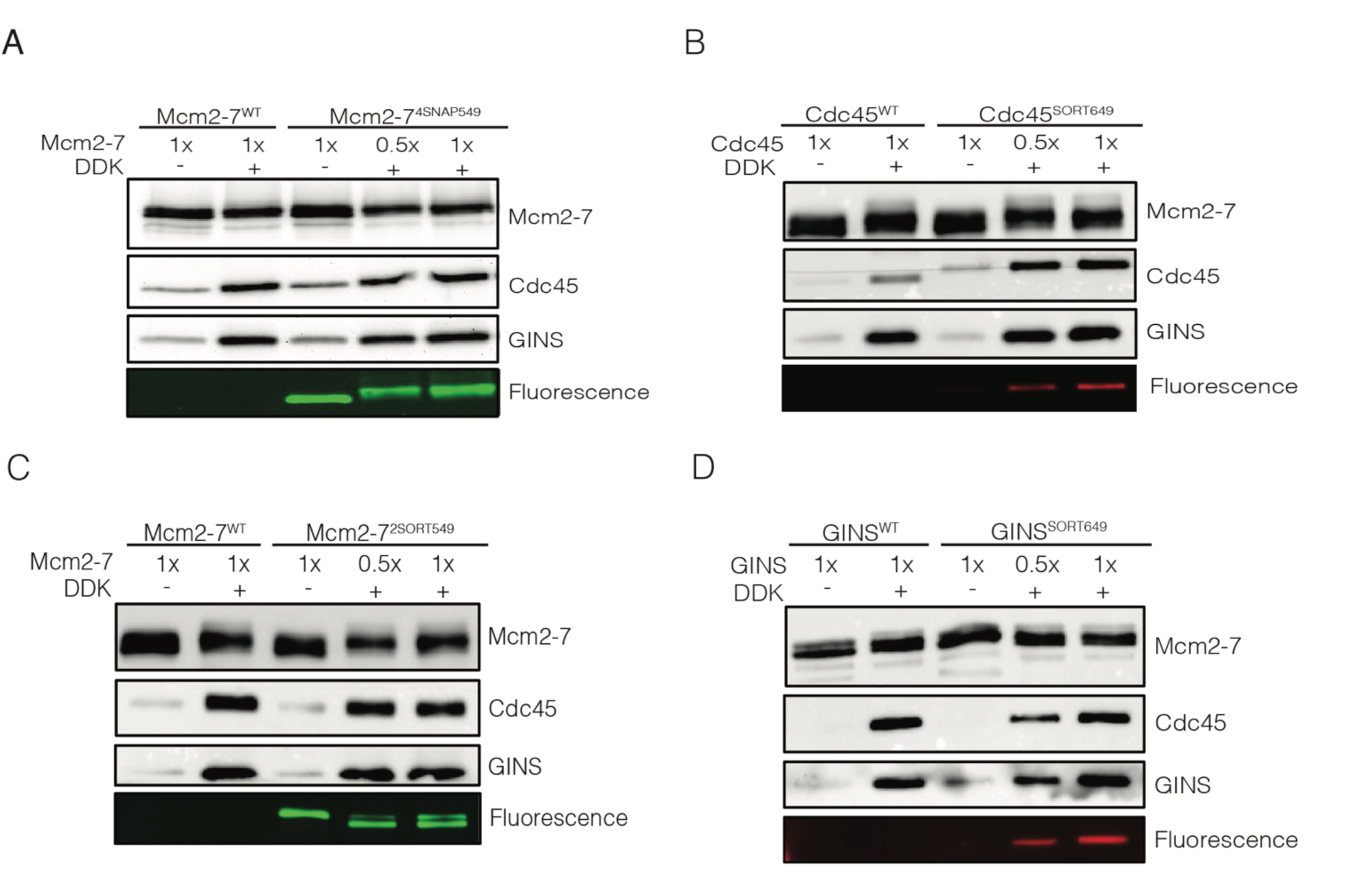
Fluorescently-labeled proteins function at wild-type levels in ensemble CMG-formation assays. Mcm2-7 was loaded onto bead-coupled 3.8-kb plasmids and subsequently phosphorylated with DDK. The phosphorylation reaction mix was removed without washing the beads prior to addition of helicase-activation proteins. Beads were washed with high-salt buffer, and Mcm2-7 proteins, Cdc45 and GINS were detected by immunoblot. Omission of DDK was used as a control for non-specific DNA binding of Cdc45 and GINS. **(A)** Mcm2-7^4SNAP549^ did not hinder Mcm2-7 loading or Cdc45/GINS recruitment. **(B)** Cdc45^SORT649^ did not hinder Cdc45 or GINS recruitment to CMG. **(C)** Mcm2-7^2SORT549^ did not hinder Mcm2-7 loading or Cdc45/GINS recruitment. **(D)** GINS^SORT649^ did not hinder Cdc45 or GINS recruitment to CMG.

**Figure 1–figure supplement 3.**
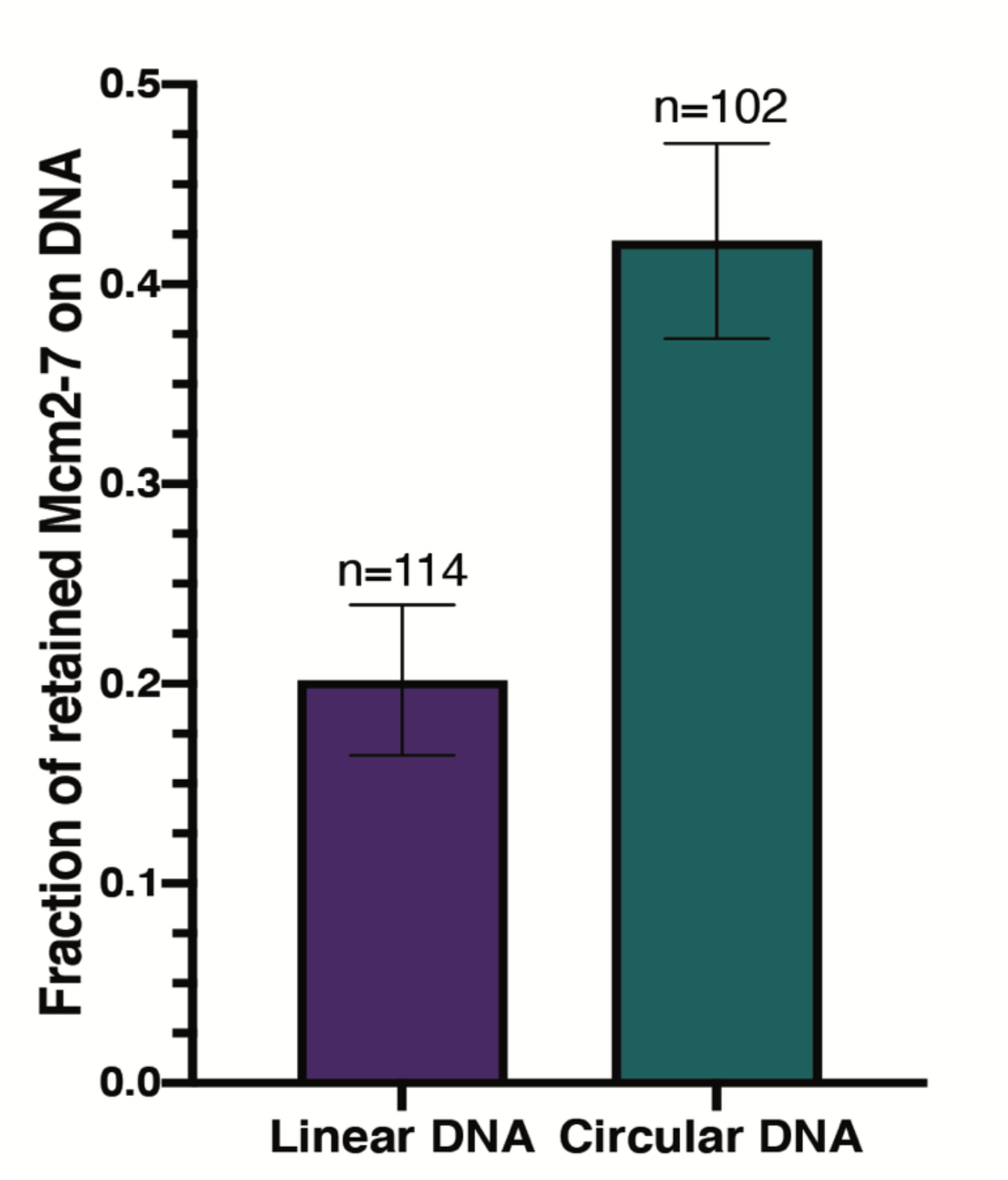
Circular DNA results in a higher number of retained Mcm2-7 during helicase loading. Bar graphs indicate the fraction (± SE) of n DNA molecules with retained Mcm2-7 during helicase loading. Origin-containing linear (purple) and circular (green) DNA are plotted.

**Figure 1–figure supplement 4.**
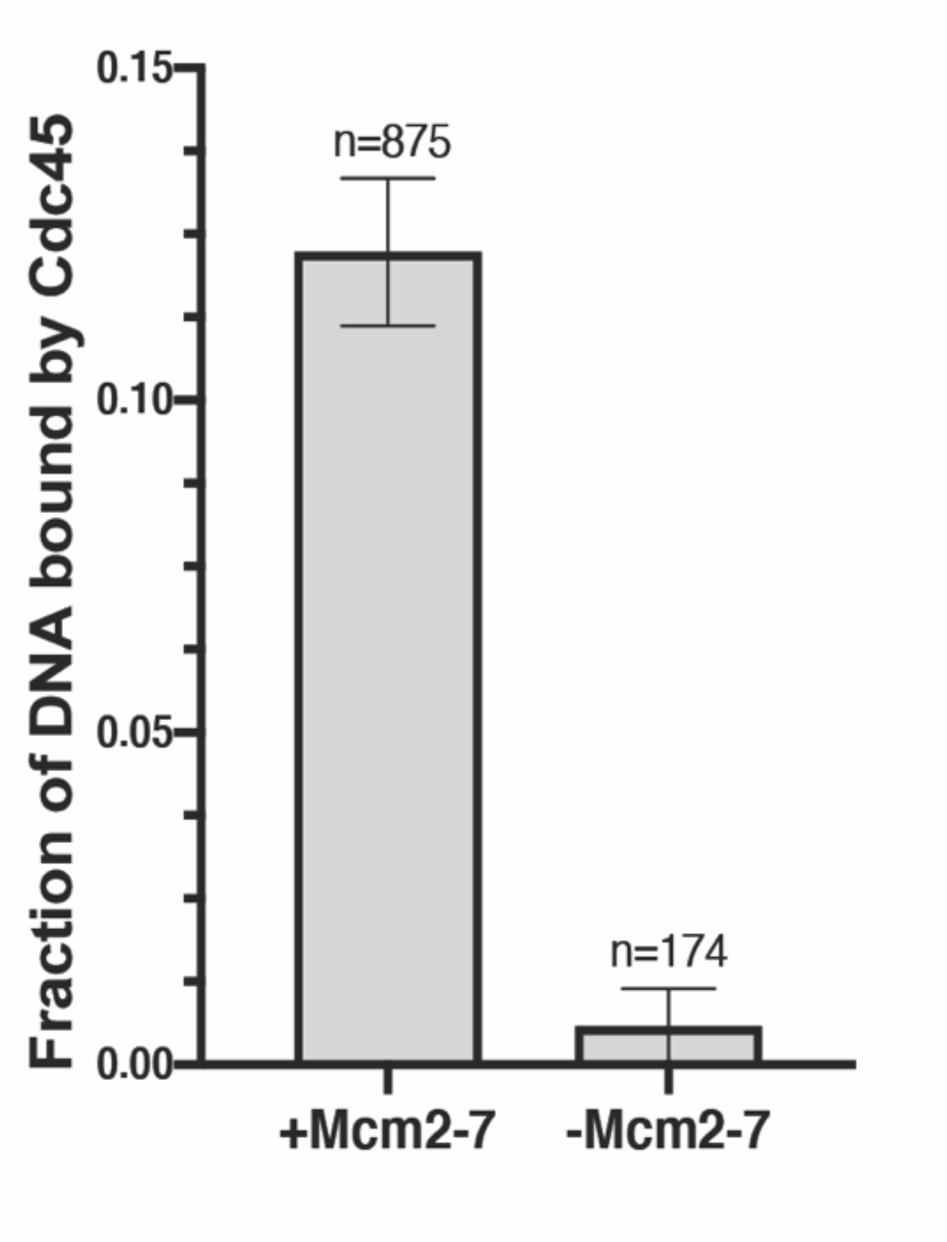
Cdc45 binds DNA in a Mcm2-7-dependent manner. The fraction of n (± SE) DNA molecules with bound Cdc45^SORT649^ after a HSW2 high-salt wash is plotted. Cdc6 was omitted from the −Mcm2-7 reaction (right bar).

**Figure 1–figure supplement 5.**
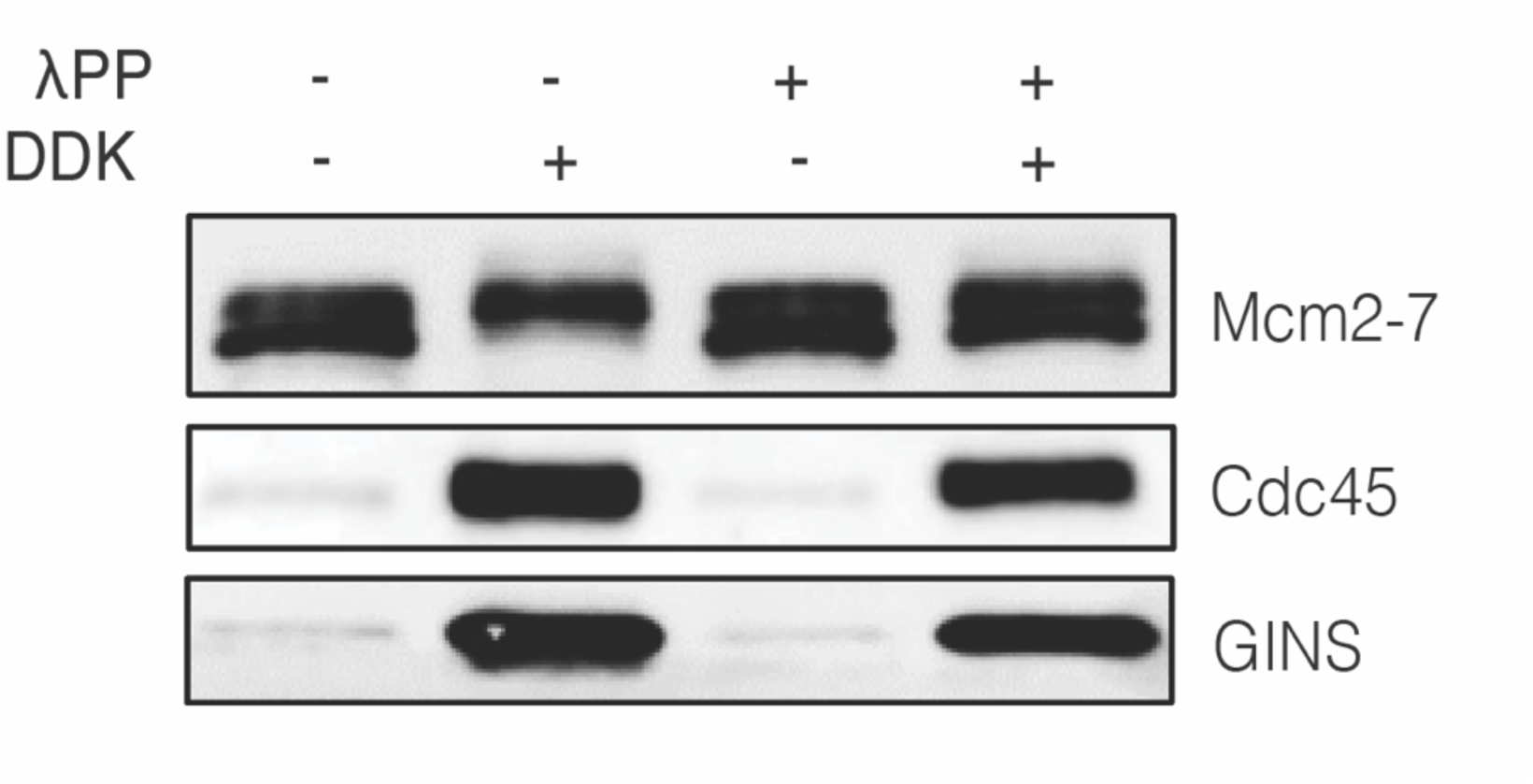
Phosphatase-treated Mcm2-7 remains functional after treatment with phosphatase. Ensemble CMG-formation assays were performed as described in Materials and Methods. Immunoblots for Mcm2-7, Cdc45 and GINS after HSW2 are shown.

**Figure 2–figure supplement 1.**
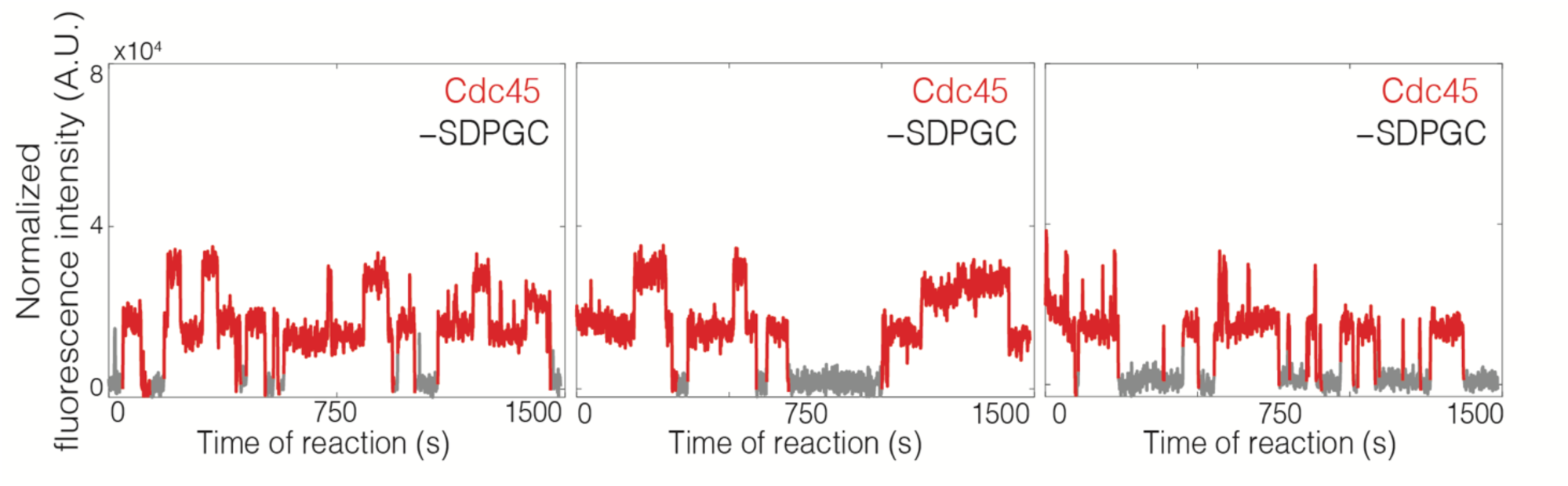
Representative records of Cdc45^SORT649^ fluorescence. Additional example traces of Cdc45^SORT649^ from the experiment shown in Figure 2A. Cdc45 recruitment reactions including Sld3/7 but without Sld2, Dpb11, Pol e, GINS, and S-CDK (SDPGC). Points colored as in Figure 2B.

**Figure 2–figure supplement 2.**
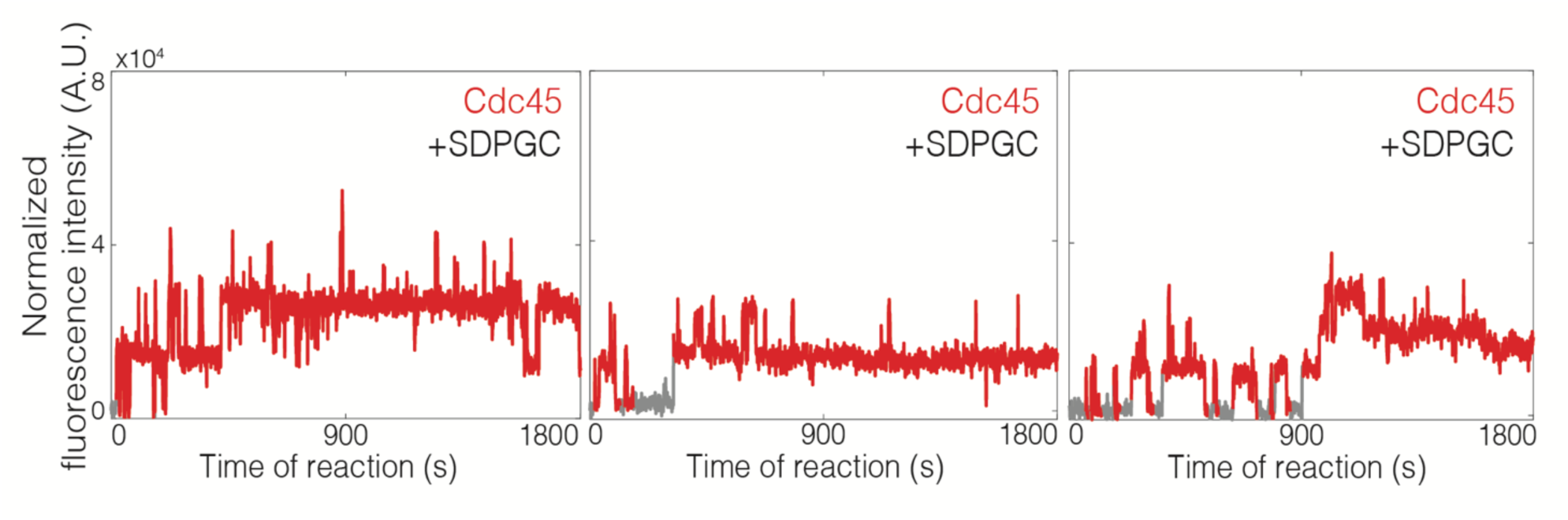
Representative traces of Cdc45^SORT649^ fluorescence. Additional example traces of Cdc45^SORT649^ from the experiment shown in Figure 2B. Cdc45 recruitment reactions including Sld3/7 in the presence of Sld2, Dpb11, Pol ε, GINS, and S-CDK (SDPGC). Points colored as in Figure 2B.

**Figure 3–figure supplement 1.**
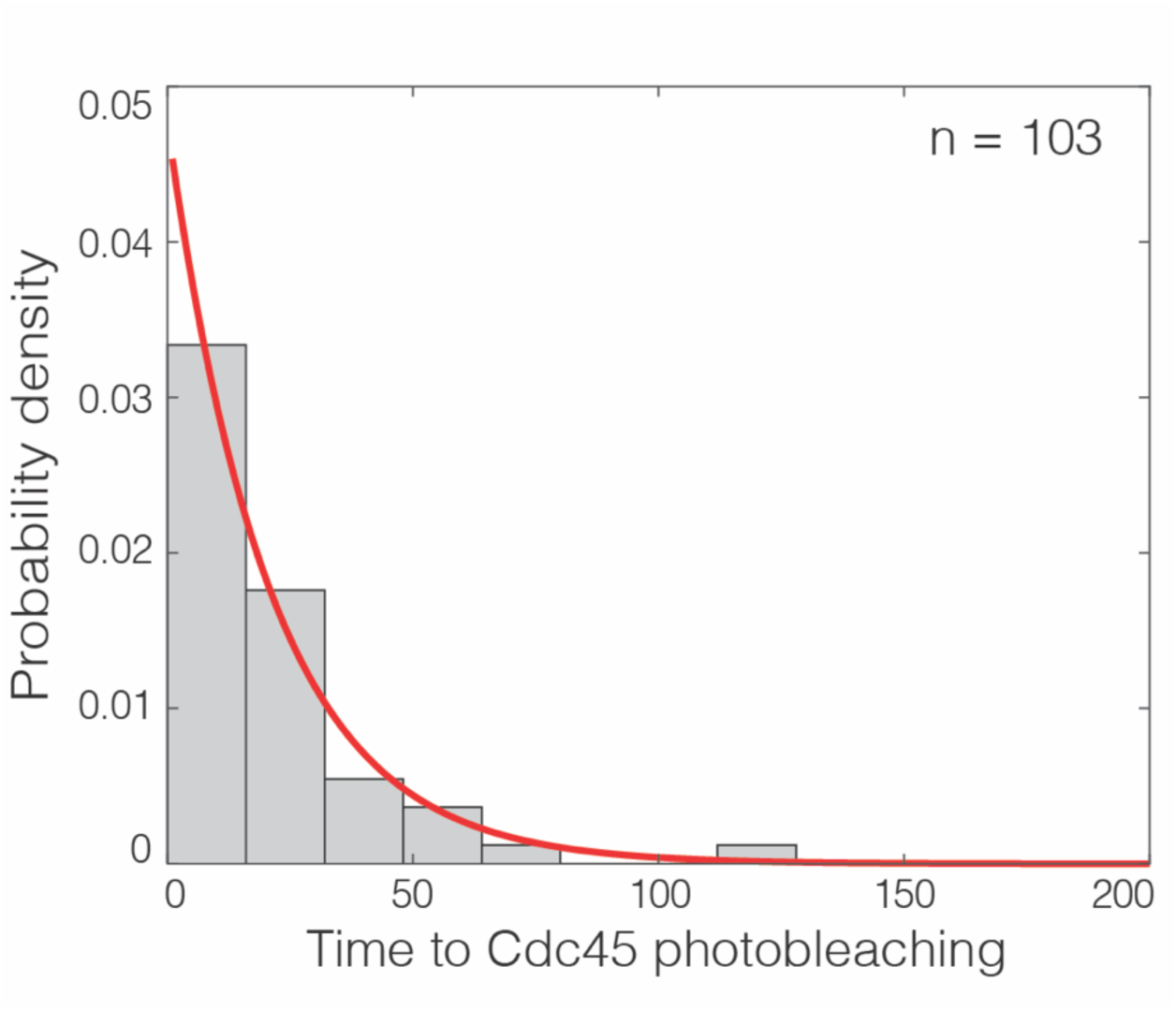
Photobleaching Cdc45^SORT649^ events are temporally resolved. Distribution of time intervals until photobleaching for Cdc45^SORT649^ bound to DNA with loaded Mcm2-7 prepared at 6.5 nM DDK, as exemplified by the lower-right record in Figure 3B. For records that showed two separate photobleaching steps, the interval was taken to be the time from the first to the second bleaching step. Exponential fit (line) yielded time constant τ = 21 s. This implies that only a small fraction (less than exp (−3 τ Δ*t*) = 7%) of sequential photobleaching steps were mistakenly counted as one rather than two steps, based on the conservative assumption all steps separated by three or more Δ*t* = 2.6 s video frame intervals were resolved.

**Figure 4–figure supplement 1.**
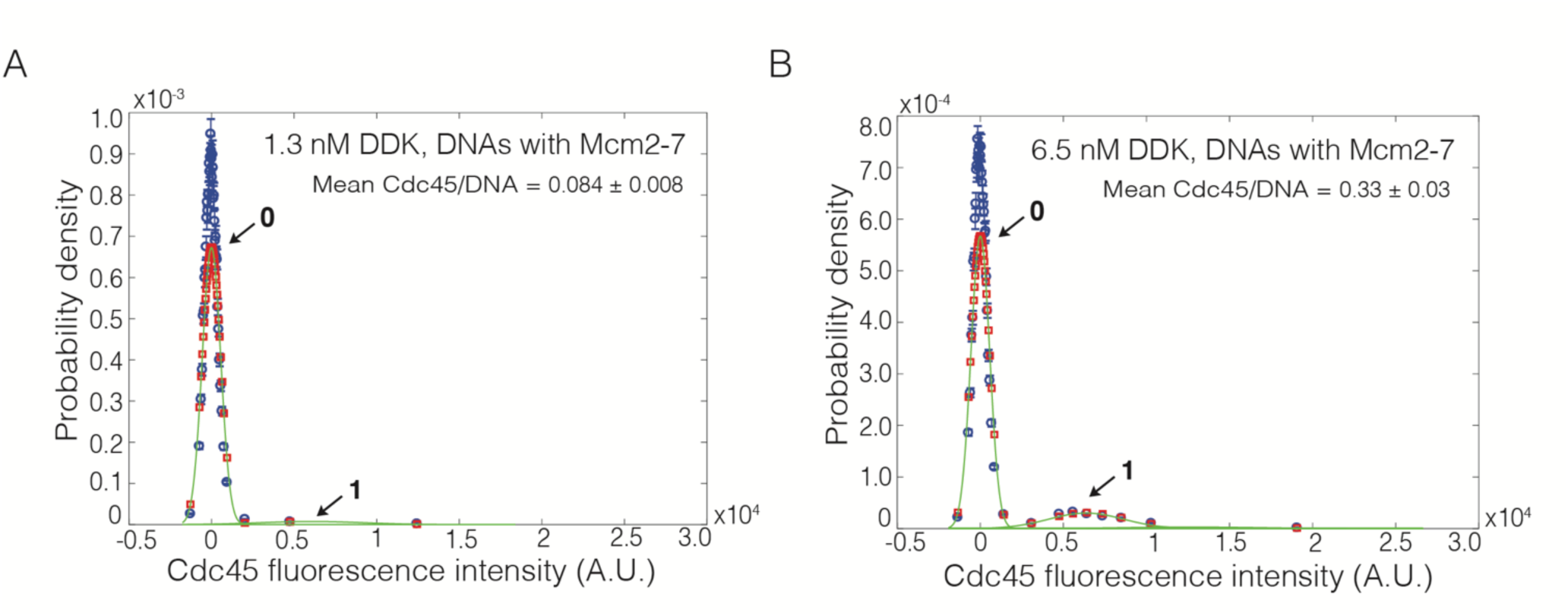
Reducing fraction of labeled Cdc45 results in detection of individual binding events. **(A and B)** CMG formation assays were performed with 3.6% of Cdc45 molecules being fluorescently labeled. Cdc45^SORT649^ fluorescence**-**intensity histograms (blue) are shown. The histogram data were fit to a sum-of-Gaussians model (red; see Materials and Methods). Fit parameters and calculated area fractions of individual Gaussian components (green) corresponding to the presence of the indicated numbers of Cdc45^SORT649^ molecules are given in Supplementary file 1 - table 1. The mean (± SE) numbers of Cdc45 molecules per DNA calculated from the fit parameters and the fraction Cdc45 labeled are indicated. **(A)** Reactions with 1.3 nM DDK on Mcm2-7-bound DNA molecules. **(B)** Reactions with 6.5 nM DDK on Mcm2-7-bound DNA molecules.

**Figure 4–figure supplement 2.**
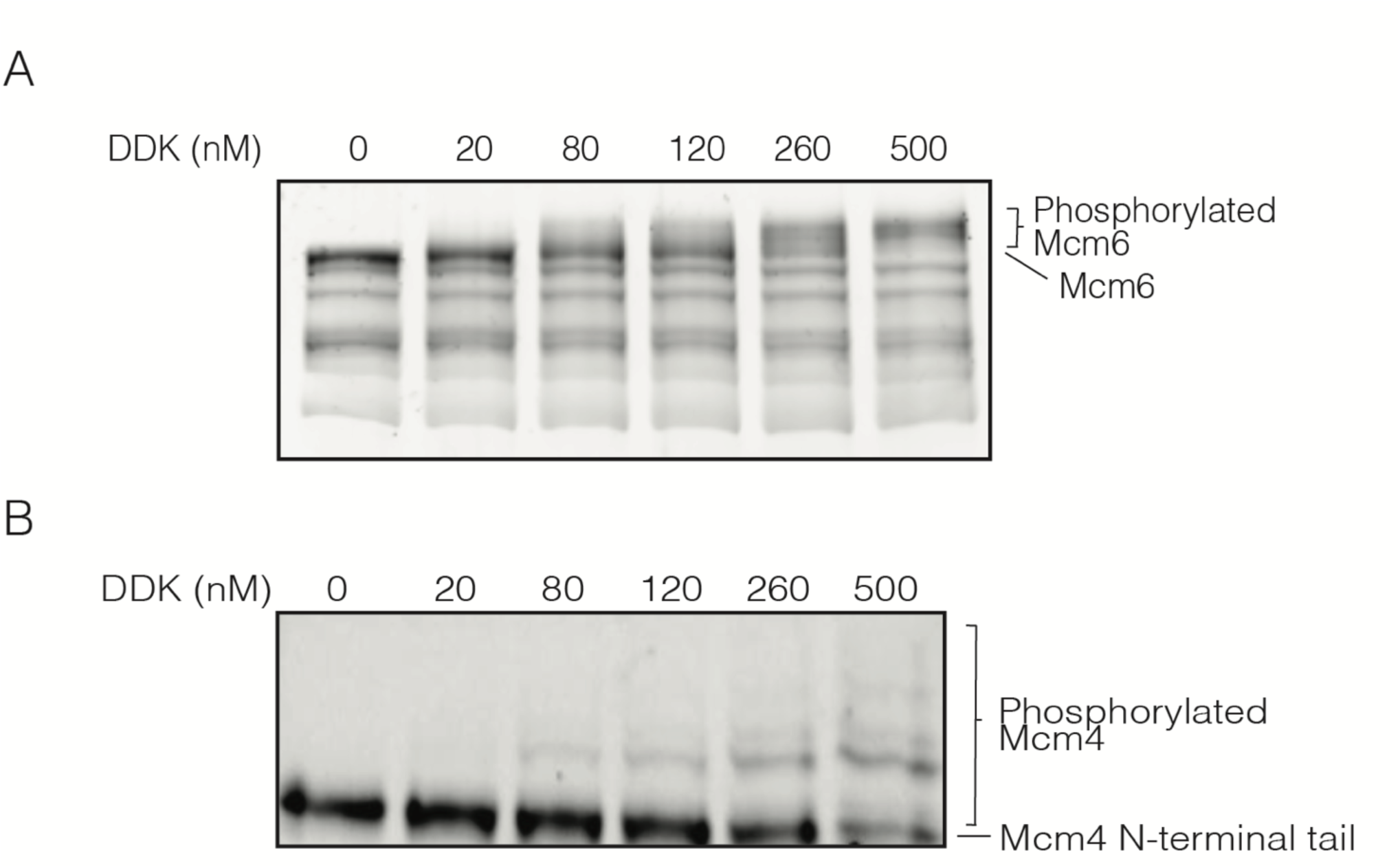
Higher DDK concentrations results in higher numbers of phosphorylation events on Mcm6 and Mcm4. **(A)** Phosphorylation of loaded WT Mcm2-7 with increasing concentrations of DDK. After loading onto DNA beads, Mcm2-7 complexes were phosphorylated with the indicated concentrations of DDK. Phosphorylated Mcm2-7 was separated on a 10% SuperSep™ Phos-tag™ gel and stained with Krypton™ Protein Stain to separate Mcm2-7 subunits with different numbers of phosphorylation sites. **(B)** Phosphorylation of Mcm4-N-terminal tail with increasing concentrations of DDK. Phosphorylated Mcm2-7 was separated on a 10% SuperSep™ Phos-tag™ gel and stained with Krypton™ Protein Stain.

**Figure 5–figure supplement 1.**
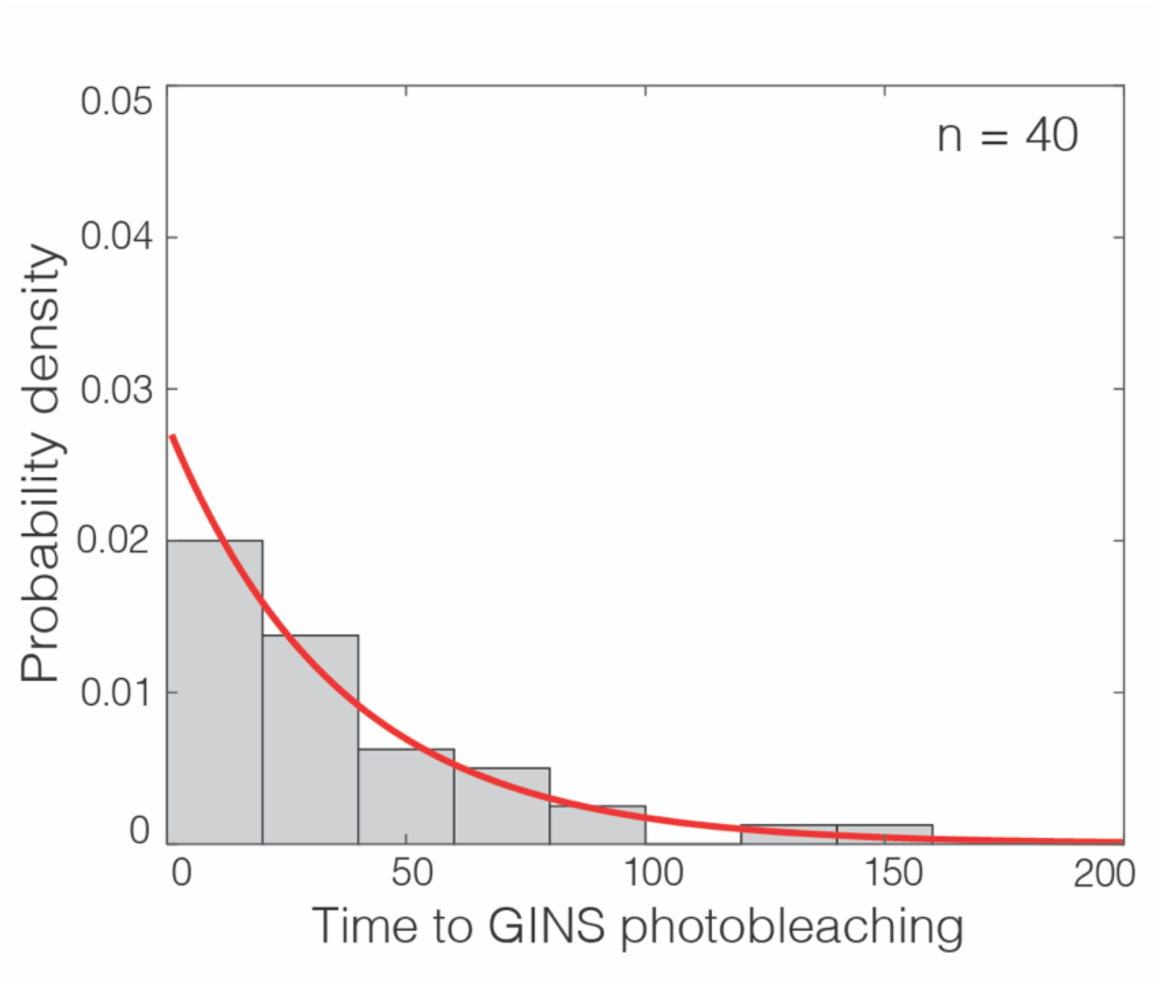
Photobleaching GINS^SORT649^ events are temporally resolved. Distribution of time intervals until photobleaching for GINS^SORT649^ bound to DNA with loaded Mcm2-7 prepared at 6.5 nM DDK, as exemplified by the lower-right record in Figure 5B. For records that showed two separate photobleaching steps, the interval was taken to be the time from the first to the second bleaching step. Exponential fit (line) yielded time constant τ = 36 s. This implies that only a small fraction (less than exp (−τ / (3Δ*t*)) = 1%) of sequential photobleaching steps were mistakenly counted as one rather than two steps, based on the conservative assumption that all steps separated by three or more Δ*t* = 2.6 s video frame intervals were resolved.

**Figure 5–figure supplement 2.**
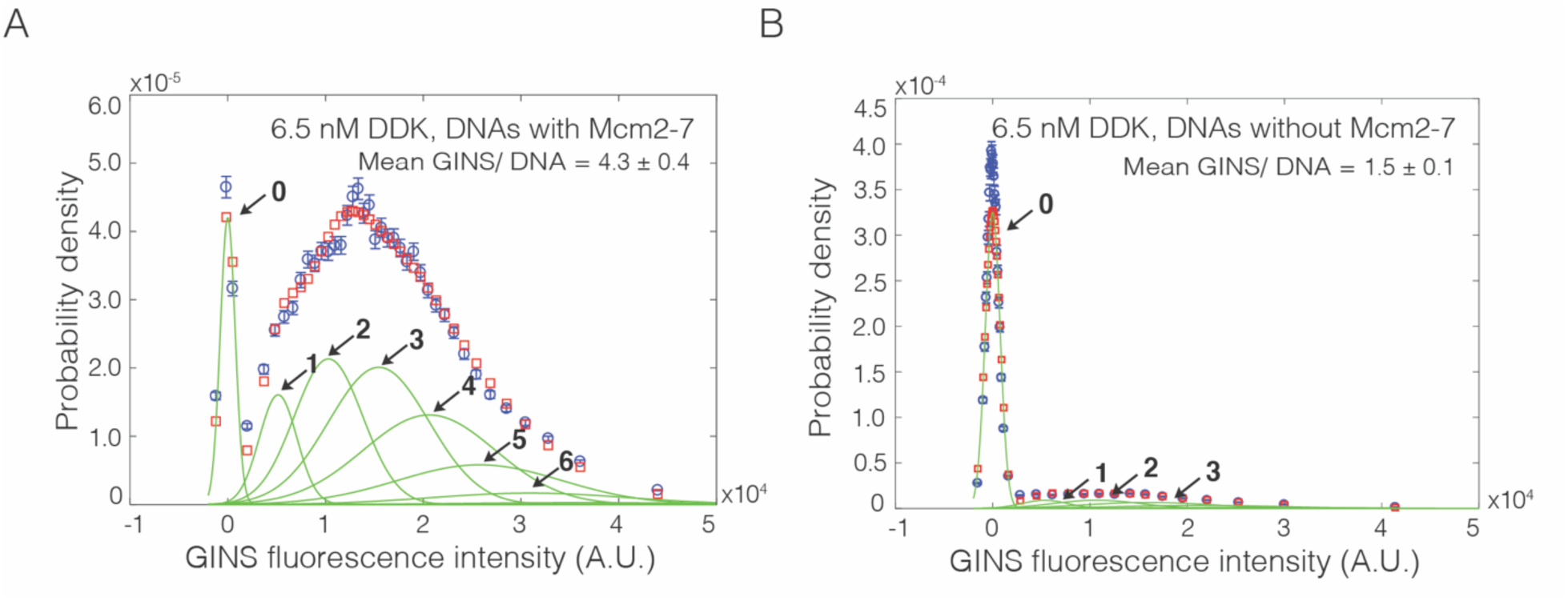
Mcm2-7^2SORT549^ dependence of GINS^SORT649^ binding. GINS^SORT649^ fluorescence**-**intensity histograms (blue) from the same recording on **(A)** Mcm2-7^2SORT549^-bound DNA (same data as Figure 5C – panel ii), and **(B)** DNA molecules without bound Mcm2-7^2SORT549^. The histogram data were fit to a sum-of-Gaussians model (red; see Materials and Methods). Fit parameters and calculated area fractions of individual Gaussian components (green) corresponding to the presence of the indicated numbers of GINS^SORT649^ molecules are given in Supplementary file 2 - table 1. The mean (± SE) numbers of GINS molecules per DNA calculated from the fit parameters and the fraction GINS labeled are indicated. Data in panel B may overestimate the amount of Mcm2-7-independent GINS binding because of the possible presence of a subpopulation of unlabeled Mcm2-7.

**Supplementary file 1–table 1.** Parameters used for fitting of Cdc45 fluorescence-intensity histograms.

| Condition<br>(Figure) | Fit parameters<sup>b</sup><br>[90% CI] |  |  |  | Fraction of time that DNAs with loaded Mcm2-7<sup>4SNAP549</sup> were bound by this<br>number of Cdc45<sup>SORT649</sup> molecules (± SE)<sup>f</sup> |  |  |  |  |  | n |
| --- | --- | --- | --- | --- | --- | --- | --- | --- | --- | --- | --- |
| | *q* | *s*<br>(× 10<sup>-3</sup>) | $\lambda$ | *x*<br>(× 10<sup>-3</sup>) | 0 | 1 | 2 | 3 | 4 | 5 | |
| Cdc45, 1.3 nM DDK,<br>–SDPGC<br>(Figure 2C – panel i) <sup>a</sup> | 0.69<br>[0.66, 0.72] | 13.41<br>[13.36, 13.49] | 0.092<br>[0.090, 0.094] | 4.12<br>[4.03, 4.21] | 0.682<br>(± 0.001) | 0.222<br>(± 0.002) | 0.079<br>(± 0.001) | 0.0159<br>(± 0.0004) | - | - | 78 |
| Cdc45, 1.3 nM DDK,<br>+SDPGC<br>(Figure 2C – panel ii) <sup>a</sup> | 0.31<br>[0.29, 0.32] | 10.10<br>[10.95, 11.05] | 0.101<br>[0.010, 0.103] | 4.19<br>[4.16, 4.23] | 0.560<br>(± 0.001) | 0.294<br>(± 0.001) | 0.116<br>(± 0.001) | 0.0262<br>(± 0.0004) | - | - | 131 |
| Cdc45, 1.3 nM DDK,<br>+SDPGC<br>(Figure 3C – panel i) <sup>q</sup> | 0.33<br>[0.30, 0.38] | 6.13<br>[6.00, 6.28] | 0.123<br>[0.116, 0.129] | 2.78<br>[2.69, 2.86] | 0.512<br>(± 0.004) | 0.295<br>(± 0.009) | 0.144<br>(± 0.004) | 0.040<br>(± 0.003) | - | - | 48 |
| Cdc45, 6.5 nM DDK,<br>+SDPGC<br>(Figure 3C – panel ii) <sup>q</sup> | 0.346<br>[0.340, 0.353] | 5.79<br>[5.74, 5.84] | 0.310<br>[0.306, 0.314] | 2.10<br>[2.07, 2.12] | 0.295<br>(± 0.001) | 0.137<br>(± 0.004) | 0.216<br>(± 0.004) | 0.194<br>(± 0.002) | 0.109<br>(± 0.002) | 0.039<br>(± 0.001) | 87 |
| Cdc45, 6.5 nM DDK,<br>+SDPGC, -Sld3/7<br>(Figure 3 – panel iii) <sup>q</sup> | 7<br>[4, 10] | 4.54<br>[4.39, 4.67] | 0.02<br>[0.01, 0.03] | 0.73<br>[0.60, 0.88] | 0.982<br>(± 0.001) | 0.017<br>(± 0.001) | - | - | - | - | 32 |
| 1:20 Cdc45, 1.3 nM DDK,<br>+SDPGC<br>(Figure 4 – supp 1) <sup>q</sup> | -0.96<br>[-0.98, -0.57] | 6.20<br>[6.16, 6.25] | 0.0003<br>[0.0001*, 0.0033] | 3.12<br>[3.03, 3.18] | 0.934<br>(± 0.001) | 0.059<br>(± 0.001) | - | - | - | - | 56 |
| 1:20 Cdc45, 6.5 nM DDK,<br>+SDPGC<br>(Figure 4 – supp 1) <sup>q</sup> | 0.6<br>[0.4, 0.8] | 6.37<br>[6.27, 6.49] | 0.048<br>[0.042, 0.055] | 2.05<br>[1.94, 2.18] | 0.797<br>(± 0.002) | 0.169<br>(± 0.003) | 0.030<br>(± 0.002) | - | - | - | 64 |
| Cdc45, 6.5 nM DDK,<br>+SDPGC, Mcm2-7<sup>Δ6N</sup><br>(Figure 6C) <sup>q</sup> | 0.8<br>[0.3, 0.9] | 7.85<br>[7.56, 7.89] | 0.017<br>[0.012, 0.019] | 4.628<br>[4.485, 4.632] | 0.925<br>(± 0.003) | 0.064<br>(± 0.004) | 0.010<br>(± 0.003) | - | - | - | 89 |

<sup>a</sup>Simultaneous 546 nm laser excitation and 633 nm laser excitation.
<sup>b</sup>See Materials and Methods
<sup>q</sup>1-sec 546 nm laser excitation followed by 1-sec 633 nm laser excitation.
<sup>f</sup>Values less than 1% were omitted.
\*Lower confidence interval uncertain because of minimal binding in this experiment.

**Supplementary file 2–table 1.**
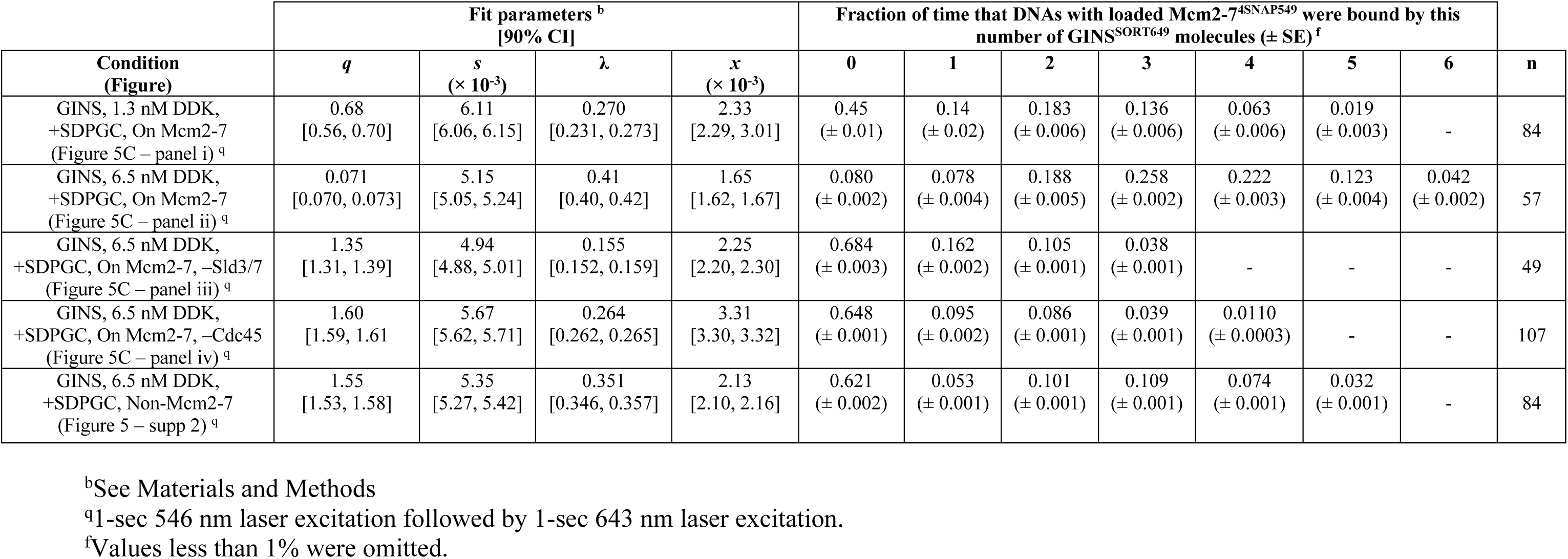
Parameters used for fitting of GINS fluorescence-intensity histograms.

**Supplementary file 3–table 1.** Yeast strains.

| Strains | Genotype | Source |
| --- | --- | --- |
| yCKR006 | his3::pSKM004 (GAL1,10-MCM2,Flag-MCM3), ura::pALS1 (GAL1,10-Cdt1, GAL4), lys::pST022 (GAL1,10-MCM4-SNAP, MCM5), trp::pCKR006 (GAL1,10-MCM6del, MCM7 | This study |
| yMM034 | ade2-1, trp1-1, leu2-3,112, his3-11,15 ura3-1, can1-100, Delta-bar1, lys2:: HisG, pep4:: unmarked, LEU:: (Cdc45-3xFlag-LPETGG) | This study |
| yST179 | yMH09 trp::pSKM003 (GAL1,10-MCM6,MCM7), ura::pALS1 (GAL1,10-Cdt1,GAL4), his::pST036 (GAL1,10-SORT-MCM2,Flag-MCM3), lys2::pSKM002 (GAL1,10-MCM4, MCM5) | Ticau et al., 2017 |
| yST147 | yMH09 trp1::pSKM003 (GAL1,10-MCM6,MCM7), his::pSKM004 (GAL1,10-MCM2,Flag-MCM3), lys2::pST022 (GAL1,10-fSNAP-MCM4, MCM5), ura::pALS1 (GAL1,10-Cdt1, GAL4) | Ticau et al., 2017 |

**Supplementary file 3–table 2.** Plasmids.

| Plasmids | Description | Source |
| --- | --- | --- |
| pCKR006 | pSK003 (GAL1,10-MCM6del, MCM7) | This study |
| pMM032 | pMM033 (Cdc45-3xFlag-LPETGG) | This study |
| pLD01 | pFJD5 (6XHIS-PreScission Protease site -Psf3 -LPETGG) | This study |

